# High-throughput single-cell CRISPRi screens stratify neurodevelopmental functions of schizophrenia-associated genes

**DOI:** 10.1101/2025.06.13.659629

**Authors:** Umut Yildiz, Annique Claringbould, Mikael Marttinen, Víctor Campos-Fornés, Mantha Lamprousi, Manu Saraswat, Mathias Saver, Daria Bunina, Michael W. Dorrity, Judith Zaugg, Kyung-Min Noh

**Affiliations:** European Molecular Biology Laboratory, Genome Biology Unit, Meyerhofstraße 1, 69115 Heidelberg, Germany; Collaboration for joint Ph.D. degree between EMBL and Heidelberg University, Faculty of Biosciences, Heidelberg, Germany; European Molecular Biology Laboratory, Molecular Systems Biology, Meyerhofstraße 1, 69115 Heidelberg, Germany; Department of Internal Medicine, Erasmus Medical Centre Rotterdam, Netherlands

**Author notes:** These authors contributed equally (Annique Claringbould, Mikael Marttinen).

## Abstract

Schizophrenia is a complex neuropsychiatric disorder with strong genetic underpinnings, yet the molecular mechanisms linking genetic risk to disrupted brain development remain poorly understood. Transcription factors (TFs) and chromatin regulators (CRs) are increasingly implicated in neuropsychiatric disorders, where their dysregulation may disrupt neurodevelopmental programs. Despite this, systematic functional interrogation in human models has been limited. Here, we combine pooled CRISPR interference (CRISPRi) screens with high-throughput single-cell multiomic profiling in hiPSC-derived neural progenitors and neurons to functionally assess 65 schizophrenia-associated genes. Based on public datasets and literature review, we selected 55 TFs and CRs, along with ten additional risk genes whose loss-of-function has been linked to schizophrenia. Our single-cell CRISPRi readouts revealed that perturbations in TFs and CRs converge on disrupting neurodevelopmental timing. CRISPRi of several factors delayed neural differentiation, whereas others, such as the knockdown of MCRS1, drove precocious neural commitment. Validation screens combined with cell cycle and metabolic indicators confirmed the differentiation-restricting or -promoting roles of these TFs and CRs.

Multimodal trajectory analysis uncovered discrete transcriptional and epigenomic states representing delayed and accelerated neurodevelopment, enriched for schizophrenia GWAS loci and disease-relevant pathways. Gene regulatory network (GRN) inference identified TCF4 and ZEB1 as critical mediators opposing the neural differentiation trajectory. Functional overexpression of these TFs followed by chromatin profiling demonstrated that TCF4 restrains, while ZEB1 promotes, neural differentiation in a stage-specific and competitive manner. Furthermore, we show that MCRS1 represses *ZEB1* expression, positioning MCRS1 as a key brake on premature neurodevelopment.

Together, our study establishes a scalable framework that integrates genetic perturbation, single-cell multiomics, and GRN modeling to functionally annotate disease-linked genes. We reveal convergent regulatory axes that underlie altered neurodevelopmental timing in schizophrenia, offering mechanistic insights into how chromatin misregulation contributes to disease pathogenesis.

## Introduction

Population-based studies have indicated a considerable genetic component underlying schizophrenia risk^1,2^, with estimated heritability rates reaching up to 80 %^2^. Although the precise etiology remains unclear, mounting evidence suggests a neurodevelopmental origin of the disease, in which genetic risk variants interfere with neurodevelopmental processes by impacting the activity of chromatin regulators (CRs) and transcription factors (TFs) that regulate cell proliferation, cell type specification, and developmental timing^3–5^. Understanding how these regulatory mechanisms function during neurodevelopment is critical to decipher the molecular underpinnings of schizophrenia. Additionally, characterizing the function of CRs and TFs in these processes can uncover their role in normal brain development and their contribution to disease states.

Pooled genetic perturbation screens have provided profound insights into gene function in proliferative cell types, including cancer and stem cells^6–12^. Such screens are powerful in annotating gene function, identifying context-specific vulnerabilities, and revealing regulatory mechanisms of selected pathways. However, gene function is highly dependent on tissue context, and therefore, it is essential to systematically interrogate gene function in disease-relevant specialized cell types and during dynamic processes like cellular differentiation. This includes neurodevelopmental processes, which are of particular disease relevance for psychiatric disorders^13^. However, primary brain material is scarce and typically only available post-mortem. Human induced pluripotent stem cells (hiPSCs) in combination with directed differentiation protocols offer a scalable and tractable model for generating homogenous populations of disease-relevant neuronal cells^14,15^. Importantly, growing evidence suggests that cortical excitatory neurons are the most affected brain cell type in schizophrenia^16–18^.

Despite advances in understanding the genetic architecture of schizophrenia^19–21^, the molecular and regulatory mechanisms of risk variants remain largely unresolved. To address this, we conducted pooled CRISPRi screens targeting schizophrenia-associated CRs and TFs in hiPSC-derived neuronal cells, coupled with high-throughput multimodal single-cell sequencing. We specifically focused on risk genes associated with loss-of-function in schizophrenia patients due to pathogenic relevance of rare coding mutations, which result in haploinsufficiency^22^. Our study revealed that CRISPRi perturbations of schizophrenia-linked CRs and TFs disrupt neural differentiation kinetics *in vitro*. We identify the convergent function of CRs and TFs in promoting neural differentiation and associate developmental pathways, including Wnt signaling, NRSF/REST, and epithelial-mesenchymal transition (EMT) with their function. In addition, we identify accelerated neurodevelopment as a potentially disease-relevant phenotype and reveal the function of MCRS1 in controlling the timing of neural differentiation with implications for brain development *in vivo*. Finally, performing gene regulatory network (GRN) interference, we demonstrate that altered neurodevelopmental trajectories are marked by the aberrant activity of specific TFs, including TCF4 and ZEB1, and their cell state-specific activities.

## Results

### Establishing high-throughput screening platforms to study schizophrenia-linked genes

We investigated the roles of loss-of-function risk genes associated with schizophrenia, focusing on CRs and TFs. By analyzing PsychENCODE datasets^23–29^ and conducting a comprehensive literature review, we identified 55 TFs and CRs (**Fig. 1a**) and seven non-TF/CR risk genes (*DGCR8*^30^, *PCCB*^31^, *MSRA*^31^, *RPTOR*^32^, *SF3B1*^32^, *SLC45A1*^32^, *THOC7*^24^) that exhibited reduced expression or activity in post-mortem brain samples from schizophrenia-affected individuals (see **Methods** for details). Additionally, we selected three well-established loss-of-function schizophrenia risk genes (*DISC1*^33^, *SETD1A*^22^, *NRXN1*^34^), resulting in a total of 65 candidate genes (targets listed in **Supplementary Data File 1**). To perturb these genes, we constructed a focused CRISPRi library consisting of 275 guide RNAs (gRNAs, four per target gene, plus 15 non-targeting controls (NT-ctrl)). The gRNA pool was cloned into a modified CROP-seq lentiviral plasmid (see **Methods** for details, gRNA sequences listed in **Supplementary Data File 1**), and gRNA complexity and distribution were verified by next-generation sequencing (area under the curve (AUC) = 0.582; all gRNAs recovered, 95 % of the gRNAs within a 3-fold distribution; skew ratio = 2; **Extended Data** Fig. 1a).

**Fig. 1.**
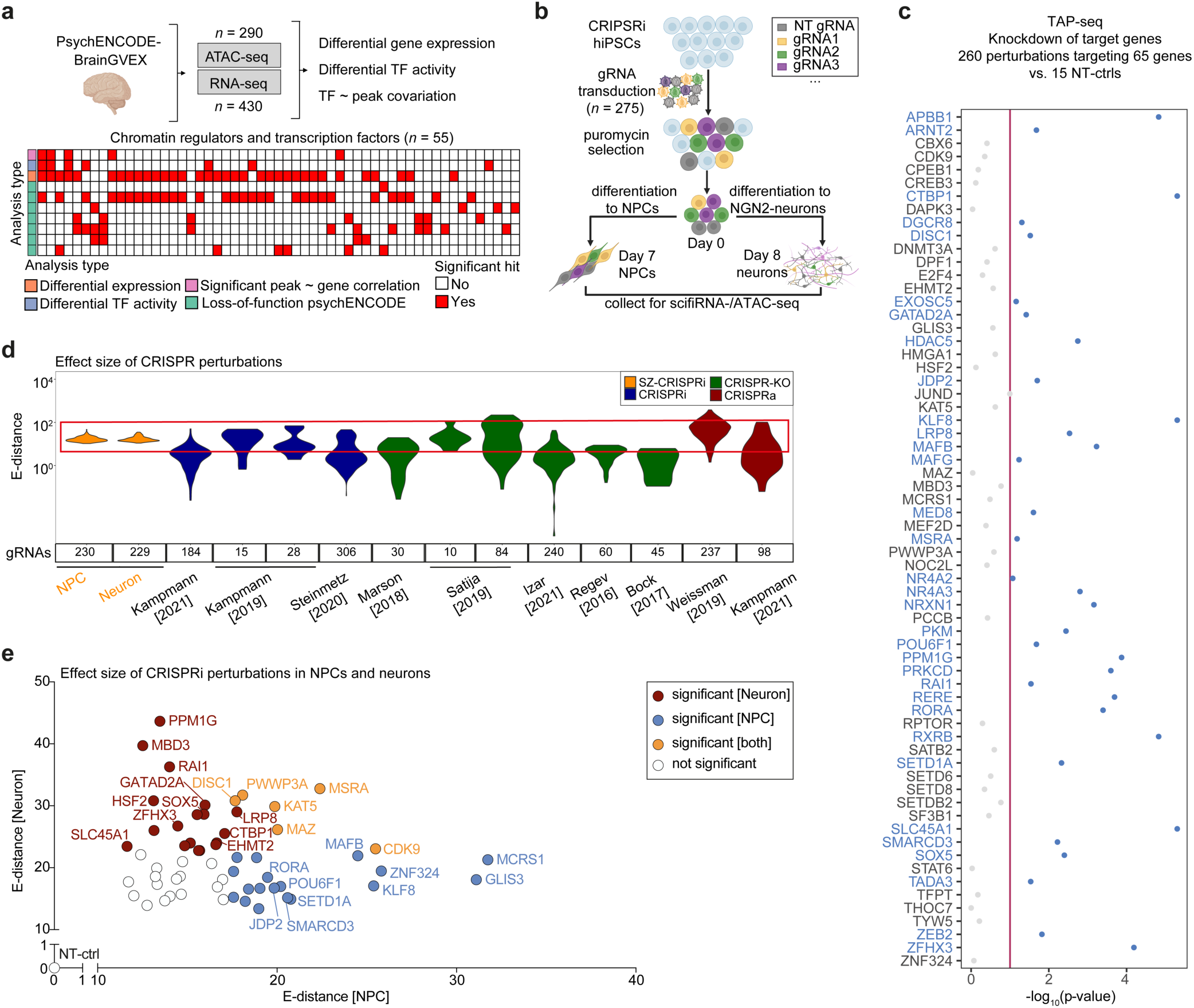
Pooled CRISPRi screening of schizophrenia-linked genes in hiPSC-derived NPCs and induced neurons with high- throughput snRNA- and snATAC-seq readouts. (a) Schematic of prioritizing schizophrenia-linked TFs and CRs (*n* = 55) using PsychENCODE datasets (green) and performing complementary analysis (orange, purple, blue) on bulk RNA-seq (*n* = 430) and matched ATAC-seq (*n* = 290) samples. List of target genes was complemented with known risk genes (*DISC1*^33^, *SETD1A*^22^, *NRXN1*^34^) and non TF/CR risk genes identified through the same analysis or comprehensive literature review (*DGCR8*^30^, *PCCB*^31^, *MSRA*^31^, *RPTOR*^32^, *SF3B1*^32^, *SLC45A1*^32^, *THOC7*^24^). (b) Strategy of the scCRISPRi screen with scifi-RNA- and scifi-ATAC-seq readouts in NPCs and induced neurons. (c) Targeted amplification (TAP-seq) of transcripts derived from schizophrenia risk genes (*n* = 65) confirmed their knockdown in the scCRISPRi screen. Pre-screening showed significant repression (*P* < 0.1 indicated on the y-axis) for ∼50 % of target risk genes (indicated along the x-axis) as compared to non-targeting control gRNAs (NT-ctrls). (d) Distribution of E-distances from CRISPRi- perturbed NPCs and induced neurons (orange, filtered dataset excluded gRNAs with low representation (≤ 5 cells)) alongside E-distances from other studies using CRISPR screens with single-cell transcriptomic readouts^14,15,44,96–101^ (datasets derived from Peidli *et al*.^45^). Number of perturbations and source of the data are indicated below the violins. For visual comparison, the E-distance distribution range from our scifi-RNA-seq screen readout is highlighted by a red box. (e) Dot plot showing significant CRISPRi perturbations according to the transcriptomic impact they induce in NPCs (x-axis) and induced neurons (y-axis) in comparison to NT-ctrl. Significant perturbations and the pairwise distance for the NT-ctrl group are highlighted in blue (NPCs), red (neurons), and yellow (both). Significant perturbations showing the strongest deviation from NT-ctrl cells are labelled (red: in neurons, blue: in NPCs, yellow: in both).

To enable functional genetic loss-of-function screens in disease-relevant cell types, we engineered hiPSCs to express a highly-effective CRISPRi repressor (dCas9-KRAB-MeCP2)^35^ and introduced a doxycycline-inducible *NGN2* cassette to facilitate rapid differentiation into excitatory glutamatergic neurons^36^. The transgenes were inserted into safe-harbor loci to ensure stable expression throughout differentiation^37,38^ (**Extended Data** Fig. 1b, c). In addition to induced neurons, we differentiated engineered hiPSCs to neural progenitor cells (NPCs) using a dual SMAD inhibition (SMADi) protocol^39^. Differentiation into NPCs and induced neurons was validated by the reduced expression of pluripotency markers (e.g., *POU5F1*, *NANOG*), and the concomitant increase in NPC (e.g., *PAX6*) and neuronal (e.g., *MAP2*) markers (**Extended Data** Fig. 1d-f). All CRISPR-NGN2 hiPSC clones maintained chromosomal integrity (**Extended Data** Fig. 1g), and induced robust gene knockdown across hiPSCs, NPCs, and induced neurons (**Extended Data** Fig. 2a-c).

To assess the effect of the schizophrenia-linked CRISPRi perturbations on cell fitness, we performed a proliferation screen in hiPSCs, NPCs, and induced neurons. CRISPRi-NGN2 hiPSCs were transduced with the gRNA pool (MOI < 0.2; *n* = 3 biological replicates), selected with puromycin, and either maintained as hiPSCs or differentiated into NPCs and neurons (**Extended Data** Fig. 2d). Guide RNA distributions in day 0 hiPSCs clustered with negative controls, indicating minimal effects of CRISPRi perturbations on cell viability or proliferation at the onset of differentiation (day 0 - four days post-transduction; **Extended Data** Fig. 2e). As expected gRNAs targeting essential genes associated with schizophrenia (*NOC2L, EXOSC5, MED8, THOC7, SF3B1*)^40,41^ were depleted in NPCs and neurons, validating the ability of the screening platform to detect phenotypic consequences of target gene knockdown (**Extended Data** Fig. 2f**, Supplementary Data File 2**). Most perturbations, however, showed little impact on cell fitness, confirming the suitability of the gRNA pool for high-dimensional analyses like scRNA and scATAC sequencing (**Extended Data** Fig. 2f**, Supplementary Data File 2**).

To increase the throughput and cost-efficiency of single-cell sequencing assays, we implemented scifi-RNA-seq^42^ for profiling gene expression, and developed scifi-ATAC-seq to assay chromatin accessibility (see **Methods** for details). These methods combine plate-based pre-indexing of split-pool of cells/nuclei with droplet-based secondary barcoding, enabling combinatorial indexing and overloading of microfluidic droplets (schematic in **Extended Data** Fig. 3a). Benchmarking with HEK293T and NIH-3T3 cells demonstrated comparable data quality to standard protocols (scifi-ATAC-seq: TSS enrichment score >10; ∼5,000 fragments per cell, **Extended Data** Fig. 3b-e; scifi-RNA-seq: ∼1,000-2,000 UMIs per cell, **Extended Data** Fig. 3f, g), at ∼10-fold lower cost. Data complexity remained consistent even with multiple nuclei per droplet (**Extended Data** Fig. 3c), and chromatin accessibility peaks were enriched at expected genomic loci, such as promoters and gene bodies (e.g., *ACTB* locus, **Extended Data** Fig. 3e).

In summary, we developed a robust hiPSC platform that integrates functional genetic screens with high-throughput single-cell sequencing workflows to comprehensively study gene function and chromatin regulation in neuronal models.

### Single-cell CRISPRi screen in hiPSC-derived NPCs and neurons

We performed a single-cell CRISPRi (scCRISPRi) screen targeting 65 schizophrenia-associated genes in NPCs and induced neurons and coupled the screen with the scifi-seq readouts. CRISPRi- NGN2 hiPSCs were transduced with the gRNA pool, selected with puromycin, and differentiated into NPCs and neurons (**Fig. 1b**). We recovered single-nucleus profiles from 41,791 NPCs and 52,673 neurons using scifi-ATAC-seq and 35,316 NPCs and 35,144 neurons using scifi-RNA-seq. In both modalities, data complexity matched the benchmark experiments (scifi-RNA-seq: median 1,254 UMIs for NPCs, and 1,816 UMIs for neurons, (**Extended Data** Fig. 3h); scifi-ATAC-seq: median 4,824 fragments per cell for NPCs and 3,844 for neurons; Transcription start site (TSS) enrichment score ∼7 for NPCs and ∼8 for neurons, **Extended Data** Fig. 3i).

To assign CRISPRi perturbations to individual cells, we amplified gRNA transcripts^43^ and integrated them into the scifi-RNA-seq workflow (see **Methods** for details), recovering gRNA identities for 60 % of NPCs and 31 % of neurons (∼450 cells per target gene, ∼2,800 non-targeting control cells), consistent with previously reported rates^14,15^. Target knockdown was confirmed for ∼50 % of schizophrenia-associated genes using TAP-seq^44^ (*P* ≤ 0.1; **Fig. 1c**; primer pool listed in **Supplementary Data File 6**). Separate qPCR assays validated the knockdown for genes not captured by TAP-seq (e.g., gRNAs targeting *EHMT2*, *SATB2*, *KAT5*; **Extended Data** Fig. 2a, b), indicating that TAP-seq may underestimate knockdown efficiencies due to technical factors (e.g., primer binding efficiency) or biological variability (e.g., alternative TSS usage, target gene expression levels).

To identify perturbations inducing significant transcriptomic variation, we applied scPerturb^45^, a benchmarking tool for scCRISPRi screens with scRNA-seq readouts. Pairwise distance scores (E-distances) between CRISPRi-perturbed cells and NT-ctrl cells indicated significant transcriptomic variation, comparable to perturbation effects observed in other scCRISPR studies from the scPerturb database (median E-distances >10, **Fig. 1d**). We observed significant transcriptomic variation in 24 and 25 of our target genes in NPCs and neurons, respectively (**Fig. 1e**; FDR ≤ 0.1).

Collectively, these results validated the functionality of the CRISPRi machinery and demonstrated its effectiveness in combination with high-throughput single-cell readouts.

### Integrated analysis reveals alterations in the neural differentiation trajectory

To leverage information from both data modalities and improve the distinction of cell states^46^, we integrated single-nucleus chromatin accessibility and gene expression readouts from the scCRISPRi screen using a knowledge graph-based neural network (GLUE^47^). The resulting shared uniform manifold approximation and projection (UMAP) showed overlap between the two data modalities and revealed a clear separation of NPCs and neurons (see **Methods** for details). Clustering the joint cell embedding for each cell type separately, we identified 17 neuron and 21 NPC clusters (Leiden algorithm; **Fig. 2a**). Although most clusters did not separate within the respective cell type, three clusters exhibited distinctive patterns. The Neuron-15 cluster formed a "bridge" between NPCs and neurons, indicating a moderate delay in neural differentiation (hereafter referred to as bridge neurons). The Neuron-12 and NPC-20 clusters exhibited an inverse pattern: Neuron-12 cells overlapped with NPCs, suggesting severe delays in neurodevelopment (delayed neurons), whereas NPC-20 cells clustered with neurons (**Fig. 2b**), suggesting premature or accelerated differentiation (accelerated NPCs).

**Fig. 2.**
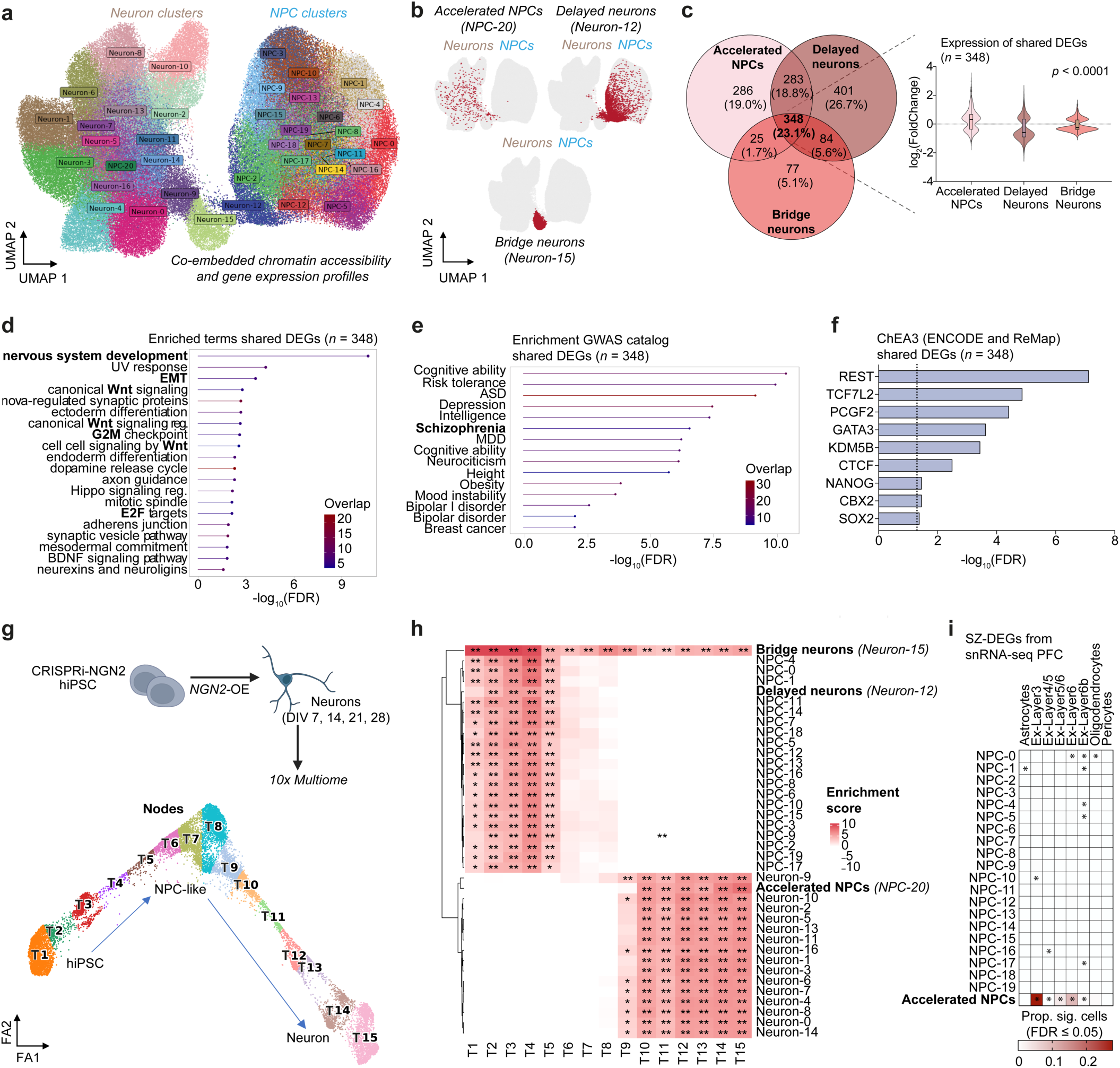
Multimodal data integration identifies cells in accelerated and delayed neurodevelopmental states. (**a**) UMAP of the co- embedded chromatin accessibility and gene expression readouts from the scCRISPRi screen (Leiden resolution: 1.2). The UMAP consists of 21 NPC, and 17 neuron clusters. (**b**) UMAPs highlighting NPC and neuron clusters associated with accelerated (NPC-20) or delayed neurodevelopmental states (Neuron-12; Neuron-15). (**c**) Overlap of DEGs in accelerated NPCs and delayed and bridge neurons. Violin plots show the expression of these genes (*n* = 348) in the three clusters. Kruskal-Wallis test was used to compute the significance score. The center line of boxes indicates the median value. Lower and upper hinges represent the 25th and 75th quantiles and the whiskers denote min/max values. (**d**) Enrichment analysis of shared DEGs with terms from the Reactome, Hallmark, KEGG, Wikipathways, and Gene Ontology databases. *EMT: epithelial-mesenchymal transition; reg.: regulation.* (**e**) Enrichment analysis of shared DEGs with traits from the GWAS catalog. *ASD: autism spectrum disorder; MDD: major depressive disorder.* (**f**) TF enrichment analysis of shared DEGs using ChEA3^103^. Dashed line indicates the significance threshold (FDR ≤ 0.05). (**g**) Top: schematic of the timecourse experiment of parental hiPSCs over the course of *NGN2*-driven differentiation. Bottom: Pseudotemporal differentiation trajectory constructed with PAGA^104^. Cells are annotated as hiPSC, NPC-like or neuron based on marker gene expression. Unsupervised clustering resulted in fifteen nodes. *FA1: forced atlas 1, FA2: forced atlas 2.* (**h**) Enrichment scores of NPC and neuron clusters from the scCRISPRi screen (y-axis) with nodes from the differentiation trajectory (x-axis). Clusters linked to accelerated or delayed neural differentiation states (NPC-20, Neuron- 12, Neuron-15) are highlighted. (**i**) Enrichment analysis using scDRS^105^ revealed significant overlap of DEGs in brain cell types of schizophrenia brain samples^18^ with DEGs derived from accelerated NPCs in the scCRISPRi screen (NPC-20 cluster). * *FDR ≥ 0.05, ** FDR ≥ 0.01* *** *FDR* ≥ 0.001.

Differentially expressed genes (DEGs; see **Methods** for details) showed significant overlap among these three clusters (23.1 % of DEGs shared between all three clusters, 41.9 % between delayed neurons and accelerated NPCs; **Fig. 2c; Supplementary Data File 3**). Notably, the shared DEGs exhibited opposing expression trends, showing elevated levels in accelerated NPCs and reduced levels in bridge and delayed neurons (**Fig. 2c**). Gene ontology analysis of shared DEGs revealed enrichment of pathways related to Wnt signaling, nervous system development, and epithelial-mesenchymal transition (EMT) (**Fig. 2d**). Furthermore, shared DEGs were significantly enriched for neuropsychiatric disorder-associated variants identified in genome-wide association studies (GWAS; **Fig. 2e**).

Transcription factor enrichment analysis of shared DEGs identified REST/NRSF, the Wnt effector TCF7L2, and Polycomb group proteins (PCGF2, CBX2) as potential effectors in the NPC-20 (accelerated NPCs) and Neuron-12/-15 (delayed/bridge neurons) clusters (**Fig. 2f)**. Motif enrichment analysis on differentially accessible chromatin regions in Neuron-12 and Neuron-15 cells confirmed the differential activity of these TFs and identified additional TFs, including TCF4, GLI3, MECOM, SOX5, EBF1, and ZEB1 (**Extended Data** Fig. 4a**, Supplementary Data File 4**). Several of these TFs have previously been associated with schizophrenia risk (e.g., TCF4^48^, ZEB1^49^, SOX5^18^) and interact with REST and Polycomb proteins (e.g., MECOM^50^, GLI3^51^, EBF1^52^, ZEB1^53^). These findings suggest their roles as key downstream effectors in neurodevelopmental processes.

To validate the accelerated and delayed developmental states in NPC-20, Neuron-12, and Neuron-15, we generated a single-nucleus chromatin accessibility and gene expression (multiome) dataset from parental NGN2-hiPSCs and induced neurons over a timecourse at days *in vitro* (DIV) 1, 2, 5, 7, 14, 21 and 28 (**Fig. 2g**). This timecourse multiome dataset revealed a differentiation trajectory composed of 15 nodes (**Fig. 2g**), with earlier nodes corresponding to progenitor-like states and later nodes aligning with differentiated neurons. By testing the enrichment of node-specific marker genes in the expression profiles of the scCRISPRi clusters (21 NPC and 17 neuron clusters), we confirmed the inverse behavior of NPC-20 (accelerated NPCs), Neuron-12 (delayed neurons), and Neuron-15 (bridge neurons), aligning with trajectory nodes corresponding to the opposite cell type (**Fig. 2h**).

To evaluate the disease relevance of these clusters, we annotated the NPC and neuron clusters using a recent snRNA-seq study of ∼500,000 nuclei from post-mortem brains of healthy and schizophrenia-affected individuals^18^, that identified cell type-specific DEGs in schizophrenia. Schizophrenia DEGs upregulated in excitatory neurons were significantly enriched in accelerated NPCs (NPC-20 cluster), but not in other NPC clusters, with up to 27 % of NPC-20 cells showing significant enrichment (FDR < 0.05; **Fig. 2i, Extended Data** Figure 4b). In contrast, delayed neurons (Neuron-12/-15) showed reduced expression of schizophrenia DEGs in excitatory neurons (**Extended Data** Fig. 4c). Overrepresentation analysis using the GWAS summary catalog^54^ further revealed that upregulated genes in accelerated NPCs were significantly associated with neurodevelopmental traits and disorders, including schizophrenia (**Extended Data** Fig. 4d).

Collectively, integrating chromatin accessibility and gene expression readouts from the scCRISPRi screen uncovered clusters of NPCs and neurons with altered differentiation kinetics.

These clusters exhibited opposing patterns in chromatin accessibility and gene expression, enriched for neurodevelopmental pathways and schizophrenia-associated genes.

### Knockdown of schizophrenia-linked genes delays neurodevelopment

The scCRISPRi screen revealed alterations in neurodevelopmental pace following the knockdown of our curated set of schizophrenia-associated genes. Given the modest effect sizes of individual genetic perturbations, which impede differential expression analysis within perturbation groups^55^, we applied two complementary strategies to link transcriptomic variation to specific CRISPRi perturbations^56,57^. First, we performed an enrichment approach^57^, comparing gRNA distributions across the defined clusters and assessing whether certain CRISPRi perturbations were overrepresented in specific clusters. Second, we grouped highly variable genes into co-expression gene modules (using Monocle3^58^, see **Methods** for details) to assess expression differences between CRISPRi perturbations.

Focusing on delayed neurodevelopmental states (Neuron-12/-15 clusters), the enrichment analysis identified 15 CRISPRi perturbations significantly overrepresented in delayed neurons (**Fig. 3a**; *P* ≤ 0.05, Fisher’s exact test). Of these, eight also showed significant transcriptional deviation from NT-control cells in the scPerturb analysis (**Fig. 1e**), supporting the hypothesis that loss-of-function in a subset of schizophrenia risk genes delays neurodevelopment^59,60^. This aligns with the concept of molecular convergence, whereby diverse genetic perturbations converge on shared downstream effects. This concept is increasingly in schizophrenia and other neurodevelopmental disorders, such as autism spectrum disorder^18,21,23,61–63^. The co-expression gene module analysis further supported the concept of molecular convergence. Nine out of the 15 CRISPRi perturbations enriched in delayed neurons showed reduced expression of neurodevelopmental gene modules (**Fig. 3b, Extended Data** Fig. 5a), which are associated with neuropsychiatric disorders (**Extended Data** Fig. 5b). Eight of these perturbations also reduced expression of neural marker genes (**Fig. 3b**), further confirming their role in delaying neural differentiation.

**Fig. 3.**
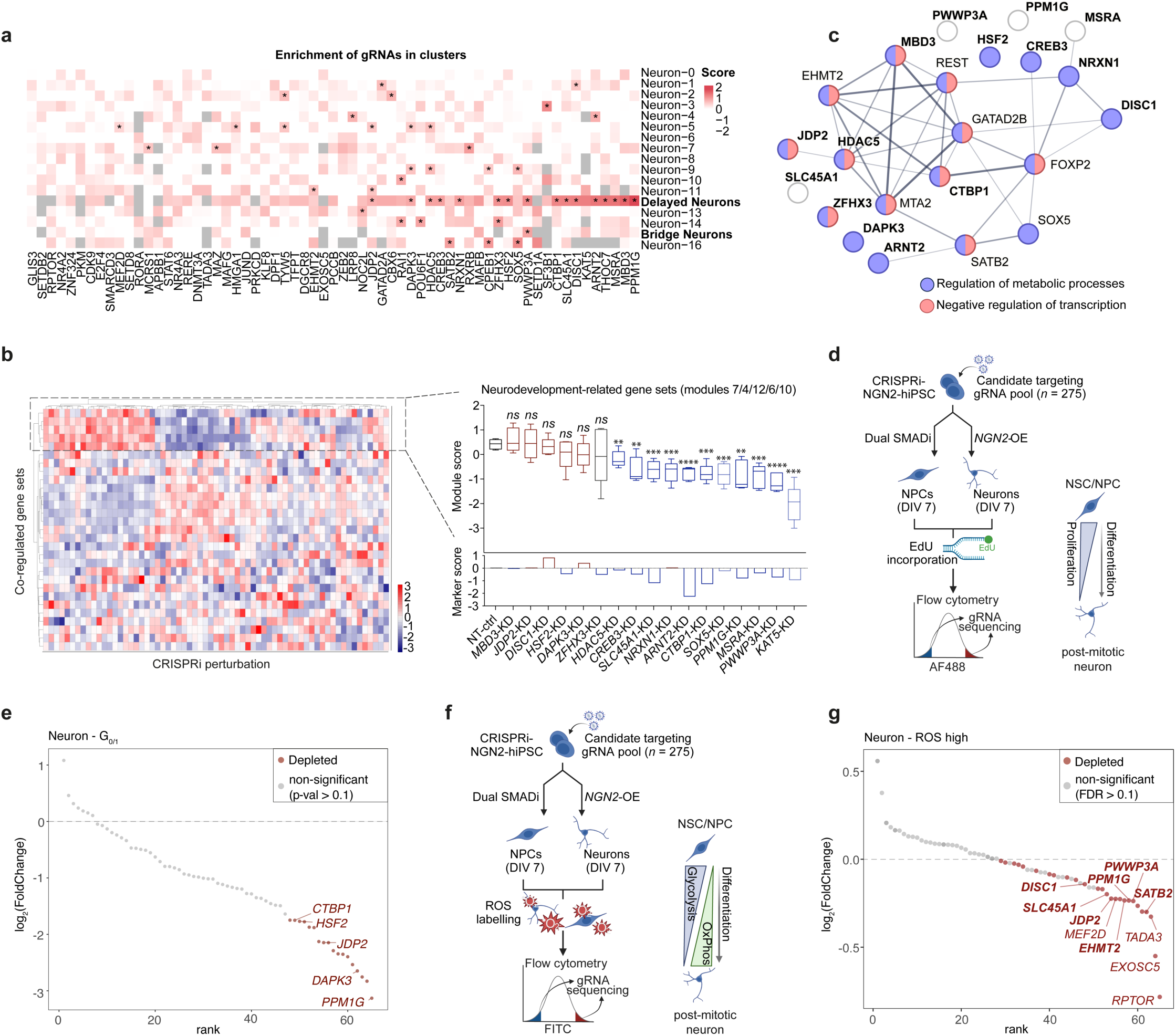
Several CRISPRi perturbations are associated with delayed developmental states in neurons, converging on common molecular mechanisms. (**a**) Distribution of gRNA identities across different neuron clusters (y-axis) collapsed on the gene level (x-axis). Significant enrichments are marked (\**P* ≤ 0.05; Fisher’s exact test). (**b**) Heatmap showing the expression of gene set modules (y-axis) following CRISPRi of schizophrenia risk genes (x-axis) in induced neurons. Monocle 3^58^ analysis retrieved 29 gene set modules. Modules 4, 6, 7, 10, and 12 (*n* = 655 genes, neurodevelopment-linked gene sets) are highlighted with a dotted box. The average module score (modules 4, 6, 7, 10, 12) contrasting the NT-ctrl group with CRISPR perturbation groups enriched in delayed neurons (**a**) shown as boxplots. The *SOX5*- and *KAT5*-KD groups were included due to their association/trend with developmental delay in the scCRISPRi screen (light blue). *THOC7*-KD was excluded due to being common essential. The center line of boxes represents the median; the lower and upper hinges represent the 25th and 75th quartiles respectively. The whiskers represent values within 1.5x interquartile range (* *P ≤* 0.05, ** *P* ≤ 0.01, *** *P* ≤ 0.001, **** *P* ≤ 0.0001, *ns:* non-significant; unpaired two-tailed t-tests). Significant hits enriched in delayed neurons (**a**) are highlighted in blue. Bottom bar shows average gene expression score of neural marker genes derived from the timecourse experiment (*n* = 150). (**c**) STRING protein-protein-interaction network comprising fifteen perturbations identified in (**a**, in bold) to induce neurodevelopmental delay. Network was extended with missing interaction partners included in the screen (*EHMT2*, *SATB2*, *SOX5*) and not (REST, GATA2B, MTA2, FOXP2). (**d**) Strategy of the FACS-based pooled CRISPRi screen to identify regulators of cell cycle progression. (**e**) Results of the EdU-CRISPRi screen in induced neurons. Significantly depleted perturbations in the G0/1-phase are marked in red. Hits from the scCRISPRi screen associated with neurodevelopmental delay are highlighted. (**f**) Strategy of the FACS-based pooled CRISPRi screen to identify modulators of intracellular ROS accumulation. (**g**) Results of the ROS-CRISPRi screen in induced neurons showing significantly less abundant (in red) hits in the ROS^high^ population. Hits associated with inducing neurodevelopmental delay in the CRISPRi screen are highlighted.

STRINGdb analysis revealed that CRISPRi perturbations enriched in the delayed neuron cluster formed predicted protein-protein interaction networks implicated in transcriptional repression and cell metabolic processes. Components of the network included HDAC5, JDP2, MBD3, CTBP1, ARNT2, and MSRA as well as known schizophrenia risk genes like DISC1 and NRXN1 that we included as additional controls (**Fig. 3c**). These results suggest that impaired transcriptional repression during *in vitro* neurodevelopment affects differentiation kinetics, leading to delayed neurodevelopmental cell states linked to schizophrenia.

Our scCRISPRi screen identified schizophrenia-associated genes that delay neural differentiation and impact commitment to the post-mitotic neuron state. To further probe the link between cell cycle dynamics and CRISPRi perturbations, we performed a FACS-based validation using EdU (5-Ethyl-2’-deoxyuridine), which is incorporated into newly synthesized DNA and marks cells in the S-phase. After pooled gRNA transduction in hiPSCs, cells were differentiated into NPCs and induced neurons (*n* = 2 biological replicates; **Fig. 3d**). Normally, induced neurons rapidly exit the cell cycle after *NGN2* expression, evading EdU labeling. In contrast, delayed differentiation results in more EdU incorporation and fewer cells in the G0/1-phase. Consistent with the scCRISPRi screen results, several CRISPRi perturbations associated with neurodevelopmental delay (e.g., *PPM1G*, *DAPK3*, *CTBP1*, *JDP2*, *HSF2*) exhibited increased EdU incorporation and decreased G0/1-phase abundance in induced neurons (*P* ≤ 0.1; **Fig. 3e**), indicating prolonged proliferation.

During differentiation, NPCs undergo a metabolic switch from glycolysis to oxidative phosphorylation to meet increased energy demands^64^. To assess metabolic changes, we labeled reactive oxygen species (ROS), primarily byproducts of oxidative phosphorylation, after pooled gRNA transduction in hiPSCs, followed by differentiation into NPCs and neurons (*n* = 3 biological replicates; **Fig. 3f**). Perturbations altering developmental timing may influence the metabolic state, with accelerated NPCs producing more ROS and delayed neurons producing less. Indeed, CRISPRi perturbations linked to delayed neurodevelopment (e.g., *PPM1G*, *DISC1*, *SATB2*, *EHMT2*, *PWWP3A*, *JDP2, SLC45A2*) showed reduced ROS levels in neurons, indicating decreased oxidative phosphorylation (**Fig. 3g**, FDR ≤ 0.1).

In summary, the CRISPRi perturbations identified several schizophrenia-associated genes that delay neural differentiation, possibly through their roles in transcriptional repression, cell cycle exit, and cell metabolism. Disruption of these processes delays neurodevelopment, underscoring molecular convergence as a shared mechanism underlying neurodevelopmental disorders.

### *MCRS1* knockdown accelerates neural differentiation

We applied the enrichment approach to the NPC readouts to identify perturbations linked to accelerated neurodevelopment. Notably, only CRISPRi of *MCRS1* was significantly enriched in the accelerated NPC cluster (NPC-20, *P* = 0.0048, Fisher’s exact test; **Fig. 4a**). *MCRS1* encodes Microspherule protein 1, a multifunctional protein involved in transcriptional regulation through the NSL (non-synthetically lethal) histone acetyltransferase complex (mediating H4K16ac deposition), the INO80 chromatin remodeling complex, and co-localization with transcriptional repressors such as HDACs^65^. Furthermore, schizophrenia GWAS identified a suggestive risk variant (*rs2303305- G*^61^) located within an exon of *MCRS1*. We applied the co-expression gene module analysis to investigate the transcriptional changes induced by the *MCRS1* knockdown (KD; see **Methods** for details). The *MCRS1*-KD led to increased expression of neurodevelopment-linked gene modules (modules 14, 16, 22, and 31; **Extended Data** Fig. 6a), enriched for biological processes such as axon guidance and postsynaptic signal transmission (**Extended Data** Fig. 6b). Many of these genes are regulated by repressive TFs involved in neuroectodermal differentiation (**Extended Data** Fig. 6c), including REST. These results suggest that decreased repressive activity on neuroectodermal genes promotes the accelerated differentiation observed in NPCs, in contrast to the inferred elevated activity of these TFs in delayed neurons.

**Fig. 4.**
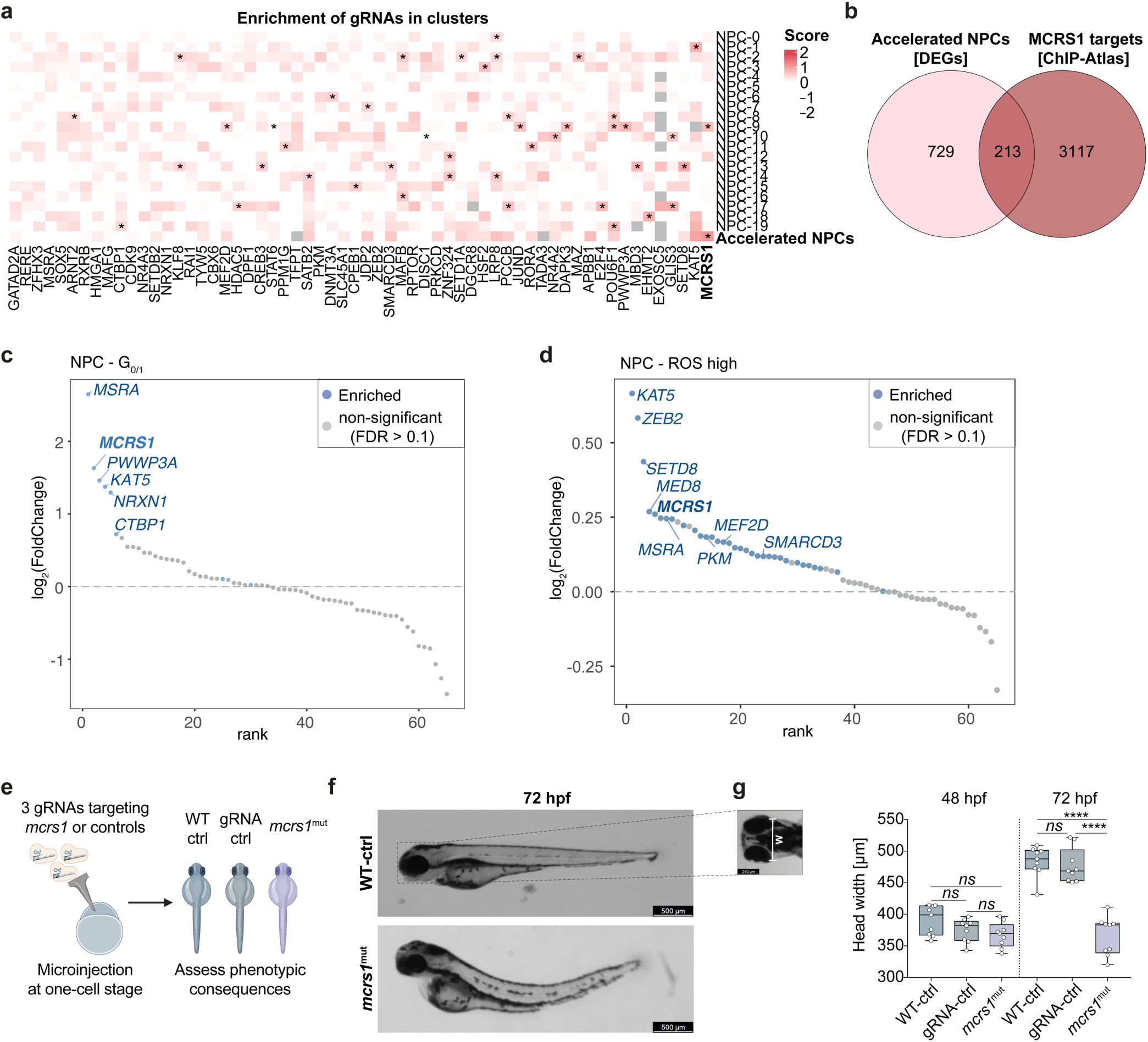
CRISPRi of *MCRS1* accelerates neural differentiation and impairs neurodevelopment *in vivo*. (**a**) Heatmap showing the abundance of gRNAs in the different NPC clusters (indicated on the y-axis). The perturbation enrichment analysis was collapsed on the gene level (indicated on the x-axis) and significant enrichments are marked (* *P* ≤ 0.05, Fisher’s exact test). (**b**) Venn diagram depicting the overlap of DEGs in accelerated NPCs (NPC-20) and MCRS1-bound genes (data retrieved from ChIP-Atlas). (**c**) Results of the EdU- CRISPRi screen in NPCs. Significantly enriched perturbations in the G0/1-phase in NPCs are marked in blue. (**d**) Results of the ROS- CRISPRi screen in NPCs showing significantly more abundant (blue) hits in the ROS^high^ population. (**e**) Schematic of the *in vivo* CRISPR mutagenesis experiment targeting *mcrs1* in F0 zebrafish embryo. (**f**) Representative brightfield images of a control wild-type zebrafish embryo (top panel) and *mcrs1* mutant zebrafish embryo (bottom panel) 72 hours post-fertilization and CRISPR editing (72 hpf). (**g**) Strategy to compare the head sizes of control and mutant zebrafish embryos (left panel). The boxplot (right panel) shows the average width of the head comparing control (WT and *slc45a2*^mut^) and *mcrs1* mutant zebrafish at 48 and 72 hpf (*n* = 9 embryos per group). The center line of the box plots represents the median; the lower and upper hinges represent the 25th and 75th quartiles respectively. The whiskers represent values within min/max. Mann-Whitney test was performed to calculate significance value. **** *P* ≤ 0.0001, *ns:* non- significant.

To further explore the genomic function of MCRS1, we utilized publicly available ChIP-seq data (ChIP-Atlas^66^) and retrieved MCRS1-bound genes. Approximately 20 % of DEGs in accelerated NPCs (213 genes) overlapped with MCRS1 binding sites (*P* < 2.2e-16, Fisher’s exact test using NPC-20 marker genes and ChIP-Atlas-derived MCRS1 targets; **Fig. 4b**), supporting a link between MCRS1 activity and the accelerated differentiation state. The overlapping genes formed a PPI network (STRINGdb; **Extended Data** Fig. 7a) and were significantly enriched for ontology terms related to epigenetic regulation of transcription, cell metabolic processes, and cell cycle regulation (**Extended Data** Fig. 7b). Many overlapping genes were downregulated in accelerated NPCs, thus the increased expression of neurodevelopmental genes may result indirectly, as a consequence of *MCRS1*-KD. In line with the predicted role of MCRS1 in cell cycle regulation and in metabolic processes, the FACS-based phenotypic CRISPRi screens (using EdU or ROS labelling, **Fig. 3d, f**) confirmed the association between *MCRS1*-KD and accelerated neurodevelopmental states. This was evidenced by the significant enrichment of *MCRS1* targeting gRNAs in the G0/1-phase (FDR = 0.0106; **Fig. 4c**) and in the ROS^high^ population within NPCs (FDR = 0.0533; **Fig. 4d**). Additional hits, such as *MSRA* and *PWWP3*, known for their roles in cell cycle regulation and DNA damage repair^67–69^ (**Fig. 4c**), and *PKM*, a key regulator of the metabolic switch from glycolysis to oxidative phosphorylation^64^, in ROS^high^ NPCs (**Fig. 4d**), further validated the readouts of the FACS-based CRISPRi screens.

Next, we compared the results of the FACS-coupled screens with transcriptional signatures of the scCRISPRi screen. Quantifying expression of S- and G2/M-phase cell cycle marker genes^70^ revealed that accelerated NPCs were transcriptionally more similar to neurons, whereas delayed and bridge neurons resembled NPC clusters (**Extended Data** Fig. 7c). To further examine metabolic changes, we analyzed the expression of glycolysis-related genes (see **Methods** for details) in the scifi-RNA-seq data. Indeed, neurons in a delayed differentiation state, like most NPCs, showed increased expression of glycolysis-related genes, while accelerated NPCs showed reduced expression, resembling neurons (**Extended Data** Fig. 7d). Notably, bridge neurons (Neuron-15 cluster) displayed a mixed phenotype in which their metabolic profile aligned with neurons, but their cell cycle signature resembled NPCs. This suggests that the metabolic transition may occur after cell cycle dysregulation in response to perturbations associated with neurodevelopment in schizophrenia.

Finally, to assess the *in vivo* relevance of MCRS1 in brain development, we performed CRISPR mutagenesis^71^ of *mcrs1* in zebrafish embryos (F0) (**Fig. 4e, Extended Data** Fig. 7e). In line with previous findings using orthogonal gene editing approaches^72^, mutant *mcrs1* zebrafish embryos displayed abnormal body shapes and significantly smaller head sizes at 72 hours post fertilization (72 hpf; **Fig. 4f, g**). These observations suggest that MCRS1-mediated accelerated neural differentiation impairs brain development *in vivo*.

In summary, we identify MCRS1 as a novel regulator of neurodevelopmental timing, restraining transcriptional reprogramming, cell metabolic shifts, and neural differentiation.

### Inferred GRN reveals reciprocal TCF4 and ZEB1 dynamics across neurodevelopment

To uncover the molecular drivers underlying accelerated and delayed neurodevelopmental states, and disrupting neural differentiation kinetics, we inferred a GRN from transcriptomic data of delayed and bridge neurons, as well as of accelerated NPCs (**Methods**). Using SCENIC^73^, which combines co-expression networks with TF motif analysis, we constructed a GRN comprising 207 TFs, 4,431 genes, and 20,171 TF-gene connections (**Fig. 5a**). Hub TFs in this network included TCF4 and ZEB1, along with PAX3, RCOR1, TEAD1, POLR3G, BACH2, SOX5, SOX6, and EZH2 suggesting their potential role as key drivers (**Supplementary Data File 5**). Among these, RCOR1 (a co-repressor of REST/NRSF), POLR3G, and ZEB1 exhibited the highest connectivity, regulating 496, 396, and 150 genes, respectively. We observed distinct regulon activity patterns in accelerated NPCs and delayed neurons compared to other cells of the respective cell type. For instance, regulon activities of BACH2 and ZEB1 were increased in accelerated NPCs, the lineage TFs SOX5 and SOX6 were prominent in bridge neurons, and regulon activities of RCOR1 and TCF4 were increased in delayed neurons (**Fig. 5b**). These distinctive regulon activity patterns corresponded to differential TF activities inferred from the scifi-ATAC-seq data (**Extended Data** Fig. 4a) and align with the established roles of these TFs in neurodevelopment. For example, *SOX5* and *SOX6* mutations are linked to neurodevelopmental disorders^60,74,75^. RCOR1 changes the balance between proliferation and differentiation in the developing brain by modulating REST activity^76^. TCF4 and ZEB1, both previously associated with schizophrenia risk^49,77,78^, play critical regulatory roles, with TCF4 proposed as a master regulator of transcriptomic alterations in schizophrenia^18,48^.

**Fig. 5.**
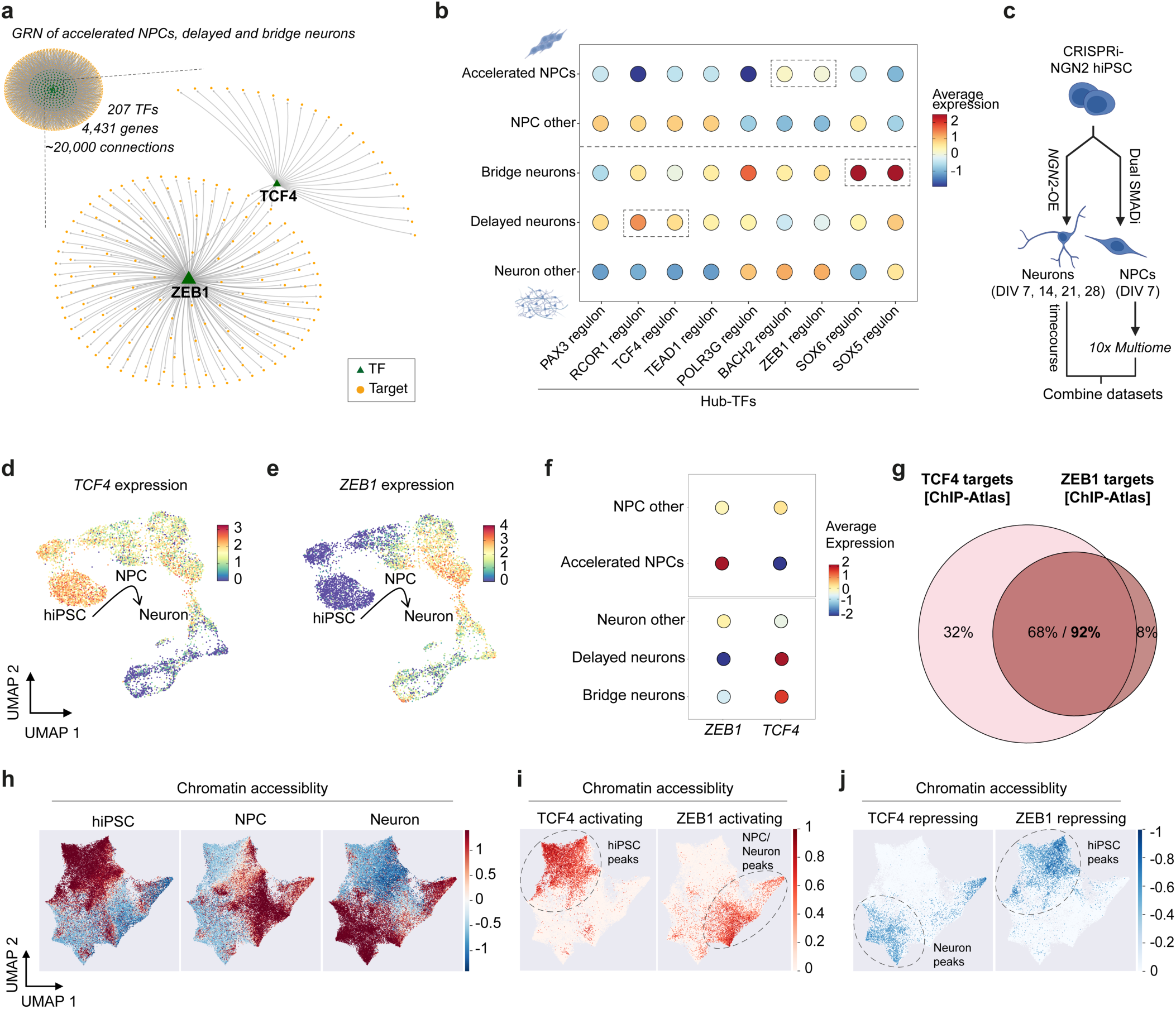
Gene regulatory network inference reveals TFs governing the accelerated and delayed neurodevelopmental cell states. (a) Gene regulatory network visualization composed of 207 TFs, 4,431 genes and 20,171 TF-gene connections. The GRN was constructed using the transcriptomic readouts of single cells of accelerated NPCs and delayed and bridge neurons. TFs in the GRN are marked with a green triangle, while target genes are indicated as yellow circles. Two of the hub-TF networks (ZEB1, TCF4) are highlighted. (b) Dot plot showing the average regulon expression of hub TFs derived from the GRN (x-axis; GRN filtered for the top 5000 connections) in accelerated NPCs in comparison to all other NPCs and delayed and bridge neurons in comparison to all other neurons (y-axis). (**c**) schematic depiction integrating timecourse multiome readouts of non-perturbed CRISPRi-NGN2 hiPSCs over the course of *NGN2*-driven differentiation with multiomic profiling of non-perturbed NPCs following dual SMADi. The NPC dataset was combined with the previous timecourse data derived from hiPSCs, and induced neurons at DIV 1, 2, 5, 7, 14, 21 and 28 post induction (**d, e**) UMAPs showing *TCF4* (**d**) and *ZEB1* (**e**) expression along the differentiation trajectory. Inferred cell states are indicated on the UMAP and each cell is colored according to *TCF4* (**d**) or *ZEB1* (**e**) expression. (**f**) Dot plots showing the average expression of *TCF4* and *ZEB1* in the NPC (top) and neuron (bottom) clusters from the scCRISPRi screen. Circles indicate the percentage of cells within each group expressing *TCF4* or *ZEB1*. (**g**) Venn diagram showing the overlap of TCF4 and ZEB1 targets (data retrieved from ChIP-Atlas of human stem and neural cell types; see **Methods** for detailed description). (**h**) scDoRI-derived UMAPs of accessible chromatin regions in hiPSCs (left), NPCs (center), and neurons (right). Analysis was performed using the NPC-embedded timecourse multiome data. (**i, j**) scDoRI-derived UMAPs showing chromatin peaks that are positively (**i**) or negatively (**j**) regulated by TCF4 (left) or ZEB1 (right).

Leveraging the multimodal differentiation timecourse data, we confirmed the central role of ZEB1 and TCF4 in NPCs and neurons through their developmental stage-specific expression and activity patterns (**Fig. 5c-e**). *TCF4* expression progressively decreased during *in vitro* neural differentiation (**Fig. 5d**), whereas *ZEB1* expression, initially low in hiPSCs, peaked at early NPCs before declining over time (**Fig. 5e**). Consistent with variation in TCF4 and ZEB1 regulon activities observed in the scCRISPRi screen (**Fig. 5b**), *TCF4* and *ZEB1* displayed opposing expression profiles in accelerated NPCs and delayed neurons compared to the respective other NPCs and neurons (**Fig. 5f**). The inverse expression and activity patterns suggest that TCF4 and ZEB1 play antagonistic roles in regulating developmental timing. Analysis of publicly available ChIP-seq datasets from brain and stem cells (ChIP-Atlas^66^, see **Methods** for details) showed an extensive overlap between ZEB1 and TCF4 binding sites (92 % of ZEB1 peaks also bound by TCF4; **Fig. 5g**), suggesting that these TFs may compete for shared binding sites such as E-box motifs in the genome, particularly in response to changes in their expression levels.

To further dissect the competitive and cell state-dependent activities and of ZEB1 and TCF4 during neural differentiation, we employed scDoRI (see **Methods** for details), a framework that links TF motif-containing chromatin regions to the expression of the corresponding TFs and of their putative target genes across the differentiation timecourse (**Fig. 5h**, **Extended Data** Fig. 8a). Consistent with the transcriptomic GRN inference, TCF4-associated chromatin regions and target genes were accessible and expressed in hiPSCs, coinciding with elevated *TCF4* expression. This positive correlation supports the role of TCF4 as a transcriptional activator at the hiPSC stage (**Fig. 5i**; **Extended Data** Fig. 8b). Contrasting this, in neurons, TCF4 motif-containing chromatin regions and target genes became accessible and were expressed only after *TCF4* expression declined, suggesting that TCF4 may repress these neuronal target genes during development (**Fig. 5j**; **Extended Data** Fig. 8d). Conversely, ZEB1 showed an opposite trend. ZEB1 motif-containing chromatin regions and target genes were open and expressed in NPCs and early neurons, aligning with elevated *ZEB1* expression and indicating an activating role of ZEB1 at these stages (**Fig. 5i**; **Extended Data** Fig. 8c). In hiPSCs, however, ZEB1 motif-associated chromatin regions and target genes were negatively correlated with *ZEB1* expression, consistent with a repressive function of ZEB1 (**Fig. 5j**; **Extended Data** Fig. 8e). These results suggest that ZEB1 can act as an activator, in addition to its known role as a transcriptional repressor, while its regulatory function is shaped by dynamic competition with TCF4 during neural differentiation.

Collectively, integrating chromatin accessibility data refined the GRN and delineated opposing roles of TCF4 and ZEB1. TCF4 restricts, while ZEB1 promotes, neural differentiation. These findings suggest that the opposing expression and activity patterns of *ZEB1* and *TCF4* in delayed neurons and accelerated NPCs have functional consequences. Specifically, elevated *ZEB1* expression in hiPSCs and NPCs, resulting from *MCRS1*-KD, may promote accelerated differentiation. In contrast, sustained TCF4 activity in NPCs and neurons, as the converging consequence of the schizophrenia-linked CRISPRi perturbations, may hinder differentiation.

### *TCF4* and *ZEB1* overexpression confirms opposing roles in neural differentiation

Our CRISPRi screen targeted genes with reduced expression in schizophrenia and therefore did not include hub-TFs identified through the GRN inference that showed increased activity, such as TCF4 and ZEB1. To assess the functional impact of increased *TCF4* and *ZEB1* expression and their activities, we performed CRISPR activation (CRISPRa) and profiled the impact on genome-wide chromatin accessibility in NPCs and neurons using bulk ATAC-seq (**Fig. 6a**). CRISPRa of *TCF4* and *ZEB1* induced overexpression (OE) of these targets and impacted the expression of the NPC identity marker *PAX6* (**Extended Data** Fig. 9a). While *TCF4*-OE increased the expression of *PAX6*, *ZEB1*-OE reduced it, suggesting opposing effects on the progenitor cell state. Overexpressing these TFs induced extensive chromatin remodeling. In NPCs, *TCF4*-OE resulted in 9,334 differentially accessible regions (DARs), while *ZEB1-*OE led to 5,144 DARs (FDR ≤ 0.1; **Extended Data** Fig. 9b). In neurons, *TCF4*-OE induced 2,330 DARs and *ZEB1*-OE induced 5,831 DARs (FDR ≤ 0.1; **Extended Data** Fig. 9c). We established a proximity-based peak-to-gene link (see **Methods** for details) using perturbation and cell type-specific DARs. Genes linked to *ZEB1*- OE induced DARs in NPCs were significantly enriched for gene sets associated with neurodevelopmental processes (**Extended Data** Fig. 9d) and neuropsychiatric traits, including schizophrenia (**Extended Data** Fig. 9e). Specifically, these genes were enriched for molecular pathways, including EMT, Wnt signaling, and Notch signaling (**Extended Data** Fig. 9d), aligning with the overrepresentation analysis of DEGs observed in the disrupted cell states in the scCRISPRi screen (**Fig. 2d**).

**Fig. 6.**
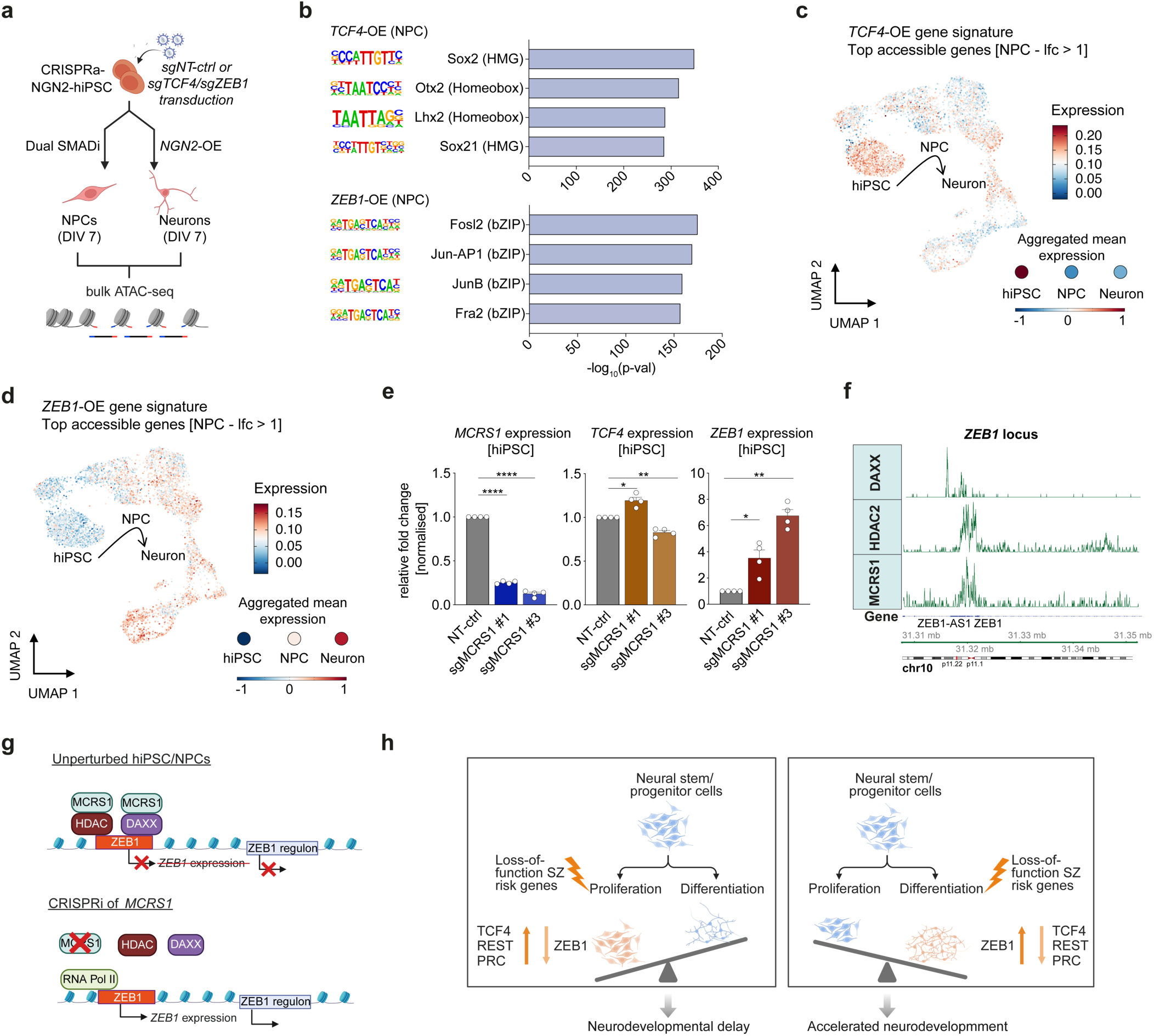
TCF4 and ZEB1 activity modulate neurodevelopmental timing. (**a**) Schematic of the bulk ATAC-seq experiment with hiPSC- derived NPCs and induced neurons following *TCF4*- or *ZEB1*-OE. CRISPRa-NGN2 hiPSCs were transduced with three gRNAs targeting the TSS of either *TCF4* or *ZEB1* or with NT-ctrl gRNAs at a low MOI (∼0.2). After selection and recovery, hiPSCs were either differentiated into NPCs (via dual SMADi) or into glutamatergic neurons (via *NGN2* overexpression). Seven days after induction, NPCs and neurons were collected, and their nuclei were isolated for subsequent ATAC-seq library preparation. (**b**) HOMER motif enrichment results in NPCs following CRISPRa of *TCF4* (top) or *ZEB1* (bottom). Top four enriched TF motifs are shown with their significance (bar plot; -log10(p-val)). (**c, d**) Expression of TCF4-OE (**c**) and ZEB1-OE (**d**) signature genes in hiPSCs, NPCs and induced neurons from the timecourse multiome experiment. Signature genes were selected using the most accessible promoter and exon regions in NPCs following *TCF4* or *ZEB1* overexpression (log2FoldChange > 1, comparison to NT-ctrl NPCs). Aggregated expression for each cell type is shown as dots. (**e**) Relative expression of *MCRS1, TCF4*, and *ZEB1* in control cells (NT-ctrl) and following CRISPRi of *MCRS1* in hiPSCs. Relative expression levels were measured via qPCR following the ΔΔCt method, normalized to *GAPDH* and *ACTB* and experimental control (NT- ctrl). Bar plots show mean ± s.e.m. (*n* = 4 independent replicates). (**f**) Genomic tracks at the *ZEB1* locus on chromosome 10 of DAXX (data retrieved from ChIP-Atlas SRX1021630), HDAC2 (ChIP-Atlas SRX19212560), and MCRS1 (ChIP-Atlas SRX3445802). (**g**) Proposed model of the regulatory function of MCRS1 on *ZEB1* expression and the associated acceleration of cell states along the neural differentiation trajectory: MCRS1 co-localizes with transcriptional repressors (e.g., DAXX, HDAC) and controls *ZEB1* expression. In this role, MCRS1 function could be considered as a “barrier”, restraining cells in the proliferative NPC state (top schematic). CRISPRi of *MCRS1* leads to the increased expression and activity of ZEB1 in hiPSCs. (**h**) Proposed model of altered neurodevelopmental trajectories and timing following the loss-of-function of schizophrenia implicated genes. Neurodevelopmental delay is the consequence of the mechanistic convergence of risk TFs and CRs in modulating the capacity of cells to repress non-neuronal transcriptional programs. On the other hand, the precocious activation of ZEB1 facilitates the fate switch and results in accelerated neurodevelopment.

To further assess how TCF4 and ZEB1 alter differentiation trajectories, we analyzed TF motifs enriched in DARs from NPCs and neurons following *TCF4*-OE or *ZEB1*-OE. DARs following *TCF4*-OE were enriched for homeobox motifs (e.g., OTX2, LHX2) and high mobility group TF motifs (e.g., SOX2, SOX21), which are linked to stemness and maintenance of the neural progenitor state^79–81^ (**Fig. 6b**, **Extended Data** Fig. 9f). In contrast, DARs following *ZEB1*-OE were enriched for pro-neurogenic bZip-TF motifs (e.g., JUN-AP1, FOSL2, FRA2)^82^, indicating a shift towards a more committed neuronal cell state (**Fig. 6b**, **Extended Data** Fig. 9f), consistent with the observed reduction in *PAX6* expression.

To connect *TCF4*-OE/*ZEB1*-OE induced chromatin changes to differentiation states, we computed gene expression scores for genes linked to differentially accessible promoter and exon regions (log2FoldChange > 1) in NPCs following CRISPRa and examined their expression across unperturbed cells during *in vitro* differentiation (**Fig. 6c, d**). TCF4-linked genes were highly expressed in hiPSCs (**Fig. 6c**), while ZEB1-linked genes showed increased expression in induced neurons (**Fig. 6d**), suggesting altered developmental states in cells overexpressing *TCF4* or *ZEB1* and highlighting the opposing regulatory function of these TFs on differentiation stage-specific gene expression.

Taken together, the GRN inference revealed dysregulated TCF4 and ZEB1 activities as key drivers of altered neural differentiation kinetics observed following CRISPRi of schizophrenia-related genes. Genome-wide chromatin profiling following the overexpression of these TFs validated their developmental stage-specific regulatory role, with TCF4 restraining neural differentiation and ZEB1 promoting it.

### MCRS1 regulates *ZEB1* expression and modulates neural differentiation pace

Importantly, the GRN inference suggested that *TCF4* and *ZEB1* expression may be directly regulated by TFs and CRs associated with delayed and accelerated neurodevelopmental state. Among these, MCRS1 targeted in our scCRISPRi screen emerged as a potential upstream regulator. To test this, we knocked down *MCRS1* and analyzed its effect on *TCF4* and *ZEB1* expression. The *MCRS1-*KD did not alter *TCF4* expression, but it significantly increased *ZEB1* expression (**Fig. 6e**). Conversely, *TCF4-*KD or *ZEB1-*KD did not affect *MCRS1* levels (**Extended Data** Fig. 9g). These results indicate a unidirectional regulatory relationship in which MCRS1 represses *ZEB1* expression. Supporting this, ChIP-Atlas data revealed MCRS1 binding peaks at the *ZEB1* promoter, overlapping with binding sites of known repressive chromatin regulators such as DAXX and HDAC2 (**Fig. 6f**), suggesting a molecular basis for the repressive role of MCRS1 on *ZEB1* (**Fig. 6g**).

In summary, the hierarchical control of *ZEB1* expression by MCRS1 underscores a regulatory axis critical for neural commitment and cell fate specification, suggesting that disruptions in this pathway during early neurodevelopment may contribute to the pathogenesis of schizophrenia.

## Discussion

Epigenetic mechanisms are central to cell type specification and the manifestation of cell identities by fine-tuning and coordinating gene expression programs. Deregulation in these processes is implicated in numerous diseases, including neurodevelopmental and psychiatric disorders. Leveraging the potential of single-cell multiomics in combination with hiPSC-based functional genetic screening, we identified several schizophrenia-associated genes, mainly TFs and CRs, with critical roles in modulating neurodevelopmental timing. Using CRISPRi, we linked the repression of these disease-implicated TFs and CRs to perturbed cell states along the neural differentiation trajectory. Chromatin regulators whose knockdown induced a delayed developmental cell state in neurons converged on common molecular mechanisms characterized by their role in transcriptional repression, including NRSF/REST and the sustained activity of TCF4. Another key finding of our work is the association of disease traits to accelerated neurodevelopment, as exemplified by the knockdown of the schizophrenia candidate risk gene *MCRS1*.

The GRN inference highlighted TCF4 and ZEB1 as key drivers of the variation observed in our scCRISPRi screen. We confirmed the central role of these TFs in both cell types by assessing their expression and activity throughout *in vitro* neurodevelopment and correlating discrepancies in their expression and activity to accelerated NPCs and delayed neurons. Genetic variation associated with increased TCF4 activity was previously linked to schizophrenia, and TCF4 was proposed to be a master regulator of gene expression differences in the disease state^18,48^. The knockdown of *MCRS1* increased *ZEB1* expression and drove NPCs to a more committed cell state, putatively by the initiation of an EMT-like program (**Fig. 6h**). In line with this, a recent study by Yan *et al.* demonstrated that ZEB1 promotes trans-differentiation of mouse fibroblasts to neurons *in vitro*^83^. Another study by He *et al*. showed that direct conversion of mouse fibroblasts to neurons requires sequential rounds of EMT and MET^84^, and Hewitt *et al*. reported accelerated *in vitro* differentiation of hiPSC lines from patients diagnosed with bipolar disorder^85^. Notably, upregulated genes in bipolar disorder-derived NPCs were significantly enriched for EMT-related genes. Importantly, precise regulation of ZEB1 activity and sequential rounds of EMT is critical not only during embryonic neurodevelopment, but also during adult neurogenesis^86^.

We validated the transcriptional consequences of our perturbations implicated in neurodevelopmental processes and neuropsychiatric disorders by phenotypic follow-up screens coupled to FACS. Lineage commitment requires precise tailoring of gene expression in a cell type-specific manner. The observed consequences of repressing schizophrenia-associated regulators in delaying neurodevelopment support the notion that cells must preserve the capacity to repress transcriptional programs related to other lineages for proper terminal commitment. Haploinsufficiency in several of these chromatin regulators (e.g., *EHMT2*, *SOX5*, *CTBP1*, *SATB2*) was previously linked to schizophrenia and other neurodevelopmental disorders, while the mechanism of action remained elusive^18,60,74,87,88^. Furthermore, neurodevelopmental timing is crucial for proper neural connectivity and function, particularly in the human prefrontal cortex, where prolonged neuronal maturation contributes to the complexity of the brain^89,90^. Neural development and maturation are closely tied to mitochondrial activity and energy metabolism of developing neurons *in vitro*^90^. In line with this, several CRs and TFs that we associated with accelerated and delayed neural differentiation impacted intracellular ROS production and cellular metabolism. Overall, our functional readouts link molecular mechanisms to the function of disease-associated chromatin and transcriptional regulators.

Our results support the hypothesis that psychiatric disorders, in particular schizophrenia, may have a developmental origin^91^. This hypothesis was first proposed by Daniel Weinberger in the 1980s^92^ and has received supporting evidence from epidemiological^93,94^ and experimental studies^61,95^. It posits that perturbations, caused by genetic and environmental factors during early brain development, are causally linked to the activation of pathological neural circuits later in life, which ultimately lead to the emergence of disease phenotypes. On the cellular level, CRISPRi perturbations altering neural differentiation kinetics may lead to immature neuronal connections that are dysfunctional and result in an overall reduced functionality of the neuronal network. This hypothesis requires future experiments examining the knockdown of *MCRS1* and NDD-linked genes on neuronal connectivity and function (e.g., via calcium imaging, multi-electrode arrays, and phagocytosis assays following co-culturing with microglia). Our CRISPR mutagenesis experiments in zebrafish embryos support this notion and suggest that accelerated neurodevelopment impairs brain development. This could be the consequence of a restricted pool of transiently amplifying progenitors that prematurely exit the cell cycle and accelerate their commitment to the neural fate, resulting in a reduced output of mature neurons.

It is important to emphasize that we associated the function of only a fraction of our selected schizophrenia-linked genes to regulatory functions in excitatory neuron development (16/65). This highlights the complexity of the disease etiology underlying schizophrenia and emphasizes the need to expand the applicability of functional genetic screens to more complex experimental model systems such as organoid, or *in vivo* screening paired with an extended set of multimodal readouts, including the spatial context, TF/CR occupancy, and proteomic changes. Nonetheless, our initial analysis on the PsychENCODE datasets derived from adult brain tissue, together with our *in vitro* screening results suggests that disease-associated TFs and CRs, whose knockdown impacted the neurodevelopmental trajectory, may play important roles not only in embryonic but also in adult neurogenesis.

In conclusion, our scCRISPRi screen provides a framework linking the function of disease-associated genes to disease-relevant neurodevelopmental phenotypes (e.g., developmental delay, accelerated differentiation), proposing putative upstream regulators of known disease-associated master TFs. By enabling systematic functional interrogation, our platform provides a foundation for generating hypotheses and guiding follow-up studies aimed at dissecting the regulatory networks underlying schizophrenia and related neurodevelopmental disorders.

## Methods

### *In-silico* selection of candidate risk genes of schizophrenia

Using metadata derived from PsychENCODE datasets^23–29^, 55 putative TFs and CRs were selected that showed an association with reduced expression and activity in post-mortem brain samples derived from schizophrenia patients. Next to the meta-analysis, differential gene expression, differential transcription factor activity and gene regulatory network analysis was performed on 430 bulk RNA-seq and 290 matched ATAC-seq samples to prioritize risk TFs and CRs, employing BrainGVEX data (Synapse ID: syn4590909). Through literature research, the list of schizophrenia-implicated genes was complemented totaling in 65 selected risk genes. The additional target genes included three positive control genes that show a strong association for disease risk (*DISC1*^33^*, SETD1A*^22^*, NRXN1*^34^) as well as seven non-TF coding risk genes (*DGCR8*^30^, *PCCB*^31^, *MSRA*^31^, *RPTOR*^32^, *SF3B1*^32^, *SLC45A1*^32^, *THOC7*^24^).

### Molecular Cloning

To introduce different dCas9 expression cassettes into the *CLYBL* donor plasmid (pC13N- iCAG.copGFP^37^, Addgene #66578), the parental plasmid was linearized via restriction digest with MluI-HF and BsrGI-HF (CutSmart, 37° C; New England Biolabs; R3198/R3575) and the resulting 10,205 bp fragment was isolated via gel electrophoresis and following the QIAquick Gel Extraction Kit (Qiagen; 28704). The linearized plasmid was ligated with PCR fragments coding for dCas9- KRAB-MeCP2^35^ (derived from Addgene #110821; FW primer: 5’- TCTGATGTACAGCCACCATGGACAAGAAGTA-3’, RV primer: 5’-ATGAACGCGTTATGAGACTCTCTCAGTCACG-3’) or dCas9-VP64^106^ (derived from Addgene #61425; FW primer: 5’-TCTGATGTACAGCCACCATGAAAAGGCCGGC-3’, RV primer: 5’-ATGAACGCGTTTACAGCATGTCCAGGTCGAAATC-3’) at a molar ratio of 1 to 4 using 400 units T4 DNA ligase (New England Biolabs; M0202) at 16° C for 16 hours. After ligation, 2 μl of the mixture was added to chemically competent bacteria (Stbl3 *E. coli*; Thermo Fisher Scientific; C7373-03) following the manufacturer’s instructions. Transformed bacteria were plated on LB-Agar plates containing Carbenicillin and incubated overnight (37° C). Small-scale liquid bacterial cultures (∼2.5 ml) were picked from the LB-Agar plates and incubated on a shaking device for approximately 10 hours (37 °C). Plasmid DNA was extracted using the Quick-DNA Miniprep Plus Kit (Qiagen; 27106) and plasmids were sequence verified (Sanger sequencing service from Eurofins Genomics) using primer flanking the insertion sites. Large bacterial liquid cultures were generated (∼250 ml) for sequence-verified bacterial clones and plasmid DNA purified following the NucleoBond® Xtra Midi EF kit (Macherey-Nagel; 740424.10).

The original pUCM-AAVS1-TO-hNGN2^38^ plasmid (Addgene #105840) was modified to exchange the puromycin resistance coding sequence for a blasticidin-S-deaminase (*BSD*) sequence. Briefly, the parental plasmid backbone was cut via restriction enzyme digestion with PmeI (New England Biolabs; R0560) and DraIII-HF (New England Biolabs; R3510) at 37 °C in CutSmart buffer. The backbone was purified via gel extraction as described above and ligated with a dsDNA gene fragment with complementary overhangs (5’- AAACGGGCCCTCTAGATTAGCCCTCCCACACATAACCAGAGGGCAGCAATTCAC GAATCCCAACTGCCGTCGGCTGTCCATCACTGTCCTTCACTATGGCTTTGATCCCA GGA TGCAGA TCGAGAAGCACCTGTCGGCACCGTCCGCAGGGGCTCAAGA TGCCC CTGTTCTCATTTCCGATCGCGACGATACAAGTCAGGTTGCCAGCTGCCGCAGCAG CAGCAGTGCCCAGCACCACGAGTTCTGCACAAGGTCCCCCAGTAAAATGATATAC ATTGACACCAGTGAAGATGCGGCCGTCGCTAGAGAGAGCTGCGCTGGCGACGCT GTAGTCTTCAGAGATGGGGATGCTGTTGATTGTAGCCGTTGCTCTTTCAATGAGGGTGGATTCTTCTTGAGACAAAGGCTTGGCCATCTCGAGCCTAGGGCCGGGATTCT CCTCCACGTCACCGCATGTTAGAAGACTTCCTCTGCCCTCTCCGCTGCCAGATCTC TCGAGGCCCTGTGGGAGGAAGAGAAGAGGTCAGAAGCTTATAACTTCGTATAAT GTATGCTATACGAAGTTATTGCCCCACTGTGGGGTGGAGGGGACAGATAAAAGT ACCCAGAACCAGAGCCACATTAACCGGCCCTGGGAATATAAGGTGGTCCCAGCT CGGGGACACAGGATCCCTGGAGGCAGCAAACATGCTGTCCTGAAGTGGACATAG GGGCCCGGGTTGGAGGAAGAAGACTAGCTGAGCTCTCGGACCCCTGGAAGATGC CATGACAGGGGGCTGGAAGAGCTAGCACAGACTAGAGAGGTAAGGGGGGTAGG GGAGCTGCCCAAA TGAAAGGAGTGAGAGGTGACCCGAA TCCACAGGAGAACGG GGTGTCCAGGCAAAGAAAGCAAGAGGATGGAGAGGTGGCTAAAGCCAGGGAGA CGGGGTACTTTGGGGTTGTCCAGAAAAACGGTGATGATGCAGGCCTACAAGAAG GGGAGGCGGGACGCAAGGGAGACATCCGTCGGAGAAGGCCATCCTAAGAAACG AGAGA TGGCACAGGCCCCAGAAGGAGAAGGAAAAGGGAACCCAGCGAGTGAAG ACGGCA TGGGGTTGGGTGAGGGAGGAGAGA TGCCCGGAGAGGACCCAGACACGGGGAGGATCCGCTCAGAGGACATCACGTG-3’) as described. Plasmid amplification was performed using chemocompetent bacteria as described. Successful cloning was verified via Sanger sequencing using sequencing primers that were spanning the editing sites.

For better estimation of the lentiviral titer, the CROP-seq-opti lentivector (Addgene # 106280) was modified via restriction digest with RsrII (New England Biolabs; R0501) and MluI-HF (CutSmart, 37° C). The linearized backbone (9,740 bp) was purified and ligated to a gene fragment coding for puromycinR-T2A-tagBFP sequence (5’-GACCGCCACA TCGAGCGGGTCACCGAGCTGCAAGAACTCTTCCTCACGCGCGTCG GGCTCGACA TCGGCAAGGTGTGGGTCGCGGACGACGGCGCCGCCGTGGCGGTCT GGACCACGCCGGAGAGCGTCGAAGCGGGGGCGGTGTTCGCCGAGA TCGGCCCGC GCA TGGCCGAGTTGAGCGGTTCCCGGCTGGCCGCGCAGCAACAGA TGGAAGGCC TCCTGGCGCCGCACCGGCCCAAGGAGCCCGCGTGGTTCCTGGCCACCGTCGGAGT CTCGCCCGACCACCAGGGCAAGGGTCTGGGCAGCGCCGTCGTGCTCCCCGGAGT GGAGGCGGCCGAGCGCGCCGGGGTGCCCGCCTTCCTGGAGACCTCCGCGCCCCG CAACCTCCCCTTCTACGAGCGGCTCGGCTTCACCGTCACCGCCGACGTCGAGGTG CCCGAAGGACCGCGCACCTGGTGCA TGACCCGCAAGCCCGGTGCCGGA TCGGGA GAGGGCAGAGGAAGTCTGCTAACATGCGGTGACGTCGAGGAGAATCCTGGCCCA CCGGTCGCCACCAGCGAGCTGATTAAGGAGAACATGCACATGAAGCTGTACATG GAGGGCACCGTGGACAACCA TCACTTCAAGTGCACA TCCGAGGGCGAAGGCAAG CCCTACGAGGGCACCCAGACCATGAGAATCAAGGTGGTCGAGGGCGGCCCTCTC CCCTTCGCCTTCGACATCCTGGCTACTAGCTTCCTCTACGGCAGCAAGACCTTCAT CAACCACACCCAGGGCA TCCCCGACTTCTTCAAGCAGTCCTTCCCTGAGGGCTTC ACATGGGAGAGAGTCACCACATACGAAGACGGGGGCGTGCTGACCGCTACCCAG GACACCAGCCTCCAGGACGGCTGCCTCATCTACAACGTCAAGATCAGAGGGGTG AACTTCACA TCCAACGGCCCTGTGA TGCAGAAGAAAACACTCGGCTGGGAGGCC TTCACCGAGACGCTGTACCCCGCTGACGGCGGCCTGGAAGGCAGAAACGACATG GCCCTGAAGCTCGTGGGCGGGAGCCATCTGATCGCAAACATCAAGACCACATAT AGATCCAAGAAACCCGCTAAGAACCTCAAGATGCCTGGCGTCTACTATGTGGACT ACAGACTGGAAAGAATCAAGGAGGCCAACAACGAGACCTACGTCGAGCAGCAC GAGGTGGCAGTGGCCAGATACTGCGACCTCCCTAGCAAACTGGGGCACAAGCTTAATTAAA-3’) incubating with T4 DNA ligase overnight (400 units, 16° C, 16 hours, 1:4 ratio of backbone to insert). Plasmids were amplified in competent bacteria, and sequence verified via Sanger sequencing as described above.

### Cloning of gRNAs targeting individual genes

For the insertion of gRNAs into the modified CROP-seq (CRISPRi) or the pXPR_502 (CRISPRa; Addgene #96923) backbone, the plasmids were linearized via BsmBI (New England Biolabs; R0580) restriction digestion (NEBuffer 3.1, 55° C), and gel purified. For oligonucleotide annealing, 100 μM of each sense and antisense oligonucleotide for a single gRNA was mixed with T4 ligation buffer (1x), nuclease-free water, and 5 units of T4 Polynucleotide Kinase (New England Biolabs; M0201). The reaction was incubated at 37° C for 45 min. Afterwards, phosphorylated oligonucleotides were annealed in the same reaction mixture by heating up to 95° C for 3 min and slow cooling to 25 °C (0.1° C per second). For the ligation, annealed oligonucleotides were diluted 200-fold and 1 μl of the dilution was mixed with 30 – 50 ng of cut plasmid backbone. After ligation, plasmid amplification and verification were performed in chemocompetent bacteria and Sanger sequencing as described above.

### Cloning of the pooled gRNA library

A focused pooled CRISPRi library was designed *in silico* targeting the 65 candidate genes with four individual gRNAs and an additional of 15 non-targeting control gRNAs using two public resources^40,107^ and ordered as an oligonucleotide pool (**Supplementary Data File 1**; Twist Bioscience). The lyophilized pool was resuspended in 3.8 μl TE buffer to yield a concentration of 10 ng/μl. In two separate reactions with 6 ng of input, the pool was amplified for ten cycles using PCR primers that annealed to the up- and downstream flanking sequences of the gRNAs (forward_primer: 5’-GGCTTTATATATCTTGTGGAAAGGACGAAACACC-3’; reverse_primer: 5’- ACTTGCTATGCTGTTTCCAGCATAGCTCTTAAAC-3’) and Q5 polymerase (PCR program: initial denaturation at 95 °C, 3 min; ten cycles of denaturation (98 °C, 20 sec), annealing (68° C, 15 sec), and extension (72° c, 15 sec) followed by a final extension at 72° C for 1 min). The modified CROP-seq backbone was cut with BsmBI, purified as described above, and additionally cleaned-up using SPRI magnetic beads (Beckman Coulter; B23318). In duplicate reactions, the amplified gRNA pool (50 ng) was cloned into the cut CROP-seq backbone (330 ng) using the NEBuilder HiFi DNA Assembly Kit (New England Biolabs; E5520) following the manufacturer’s instructions. The product was purified via SPRI magnetic beads, and the gRNA pool was amplified utilizing Endura electrocompetent cells (Lucigen; 60242-2-LU). Briefly, two electroporation reactions were performed on a MicroPulserTM II (BioRad) following the manufacturer’s instructions and using the EC1 setting (V: 1.8 kV). Subsequently, cells were resuspended in pre-warmed recovery medium (1 ml), pooled together in a 50 ml conical tube and incubated at 37° C for 45 min in a shaking-incubator. Next, the cell suspension was diluted 1:1 with LB-medium and 2 ml of bacterial suspension was spread evenly on pre-warmed LB-agar plates (245 mm square dish) containing carbenicillin (100 μg/ml) and incubated for 12 hours at 37° C. To assess the bacterial transformation efficiency, serial dilutions of the stock bacterial suspension (1:1,000; 1:10,000; 1:100,000) were plated on 10 mm LB-agar plates containing carbenicillin (100 μg/ml). Upon verification that the electroporation efficiency met the minimum recommended library representation of 100-fold bacterial colonies (∼13,000-fold representation) for each gRNA, cells were harvested with LB-medium and a cell scraper. Subsequently, bacterial cells were centrifuged (6,000x g, 20 min, 4° C) and plasmid DNA purification was performed using NucleoBond® Xtra Midi EF (Macherey-Nagel) according to the instructions provided by the manufacturer.

### Tissue culture

Human induced pluripotent stem cells with institutional review board approval (CESCG-295, kindly provided by Dr. Michael Synder, were derived from peripheral blood mononuclear cells; Stanford University, reference numbers 29904, 30064) were maintained under serum-free conditions in E8 medium (Gibco, A1517001) on vitronectin-coated (Thermo Fisher Scientific, A14700) plates at 37 °C with 5 % CO2. For passaging, cells were washed twice with phosphate-buffered saline (PBS) and incubated in a Versene solution (Gibco; 15040066; 3 min at 37 °C) to release colonies from the plate. A fraction of the cell suspension, typically 1:7 (v/v), was transferred to a freshly coated plate with E8 medium supplemented with 10 µM Rho kinase inhibitor Y-27632 (ROCKi; Abcam; Ab120129) and 100 units/ml penicillin/streptomycin (Gibco; P4458). The medium was replaced every day with fresh E8, and cells were maintained until reaching ∼80 % confluency. CRISPRi/a- NGN2-hiPSCs were maintained in E8 medium supplemented with 100 µg/ml G418 (Invivogen; Ant-gn-1) and 2.5 μg/ml blasticidin-S-hydrochloride (Sigma-Aldrich; SBR-00022). Following gRNA transduction, cells were selected with 1 to 4 μg/ml puromycin (Invivogen; Ant-pr-1).

Human embryonic kidney 293 (HEK-293T) cells and mouse fibroblast NIH-3T3 cells were cultured in DMEM, high glucose (Gibco; 11965092) supplemented with 10 % FBS (Gibco; 10270-106), 1x non-essential amino acids (NEAAs; Gibco; 11140050), 2 mM GlutaMAX (Gibco; 35050061), and 100 units/ml penicillin/streptomycin (Gibco) and maintained at 37 °C with 5 % CO2.

All cell lines were regularly tested for mycoplasma contamination using a commercial mycoplasma detection kit (E-Myco Plus PCR Detection Kit (Chembio, 25237).

### Generation of clonal hiPSC lines

Human iPSCs were maintained as described above until reaching ∼80 % confluency. Two hours before nucleofection, the cell culture medium was replaced with E8 medium supplemented with ROCKi (10 μM). For each nucleofection reaction, one million cells were collected via Accutase (Stem Cell Technologies; 07920) treatment (5 min, 37 °C) in PBS. Following centrifugation (250x g, 5 min), cells were resuspended in the transfection mixture following the manufacturer’s recommendations (Lonza; V4XP-3024). Nucleofections were performed with 10 μg of the respective transfer plasmid encoding for the transgenes (e.g., CRISPRi/a machineries, NGN2 expression cassette) and 5 μg of each TALEN-encoding plasmid with homologies towards the 5’- and 3’-ends of the target locus (*CLYBL* left: Addgene plasmid #62196, *CLYBL* right: Addgene plasmid #62197, *AAVS1* left: Addgene plasmid # 80495, *AAVS1* right: Addgene plasmid # 80496). Cells were transfected on the 4D-Nucleofector using the “CA-137” program. After electroporation, cells were gently transferred to 15 ml conical tubes with pre-warmed E8 medium and centrifuged (220x g, 5 min). The cell pellet was resuspended in pre-warmed E8 medium supplemented with ROCKi (10 μM), and cells were transferred onto vitronectin-coated plates. The cell culture medium was replaced every day, and cells were passaged as described above. Two weeks after nucleofection, the respective selection marker was added to the cell culture medium (G418, blasticidin-S-hydrochloride, or both). Following two weeks of antibiotic selection, cells were FAC- sorted onto vitronectin-coated 96-well plates to establish clonal transgenic cell lines under exclusion of dead cells via DRAQ7 (Biolegend, 424001) staining. The CRISPRi-NGN2 hiPSCs were sorted based on the expression of BFP that is fused to dCas9 and mCherry that is part of the inducible *NGN2* expression cassette, while CRISPRa-NGN2 hiPSCs were sorted based on mCherry expression only. One week after FACS, the antibiotic selection was re-initiated as described. Surviving clones were harvested and genotyped for the correct insertion of the transgenes (FW_5’_genotyping_CLYBL: 5’-GCATTCTGCTTGGGAACAACA-3’, RV_5’_genotyping_CLYBL: 5’-GATGCGATGTTTCGCTTGGT-3’; FW_3’_genotyping_CLYBL: 5’-TGTATTTGTGGACTTCACCAGGG-3’,RV_3’_genotyping_CLYBL5’-<colcnt=3> CCACCAGCAACCTGACGTTTT-3’;FW_5’_genotyping_AA VS1:CCAAAAGGCAGCCTGGTAGA-3’, RV_5’_genotyping_AAVS1:5’-TTTACGAGGGTAGGAAGTGGT-3’; FW_3’_genotyping_AAVS1:5’-TGACACCAGTGAAGATGCGG-3’,RV_3’_genotyping_AAVS1:5’-TCGACCTACTCTCTTCCGCA-3’)). Upon expansion, a bulk sort was repeated to enrich for cells with high expression of the transgenes using the same gating strategy as described for the single-cell sorting.

Maintenance of the chromosomal landscape was verified via low-coverage whole-genome sequencing as described before (Matteo’s paper). Final sample libraries were equimolarly mixed and sequenced as a multiplexed pool on the NextSeq 500 platform (Read configuration: 75 cycles read 1, 6 cycles index 1, single-end reads).

### Differentiation of hiPSCs to neural progenitor cells and induced neurons

For the differentiation of hiPSCs to NPCs, a monolayer differentiation protocol was employed following the SMADi Neural Induction Kit instructions (Stemcell Technologies; 0858). Briefly, 12- or 24-well tissue culture plates were coated with 15 μg/ml poly-L-ornithine (Sigma-Aldrich; P4957) and 10 μg/ml laminin (Roche; 11243217001) before cell collection. Next, hiPSCs were washed twice with PBS and dissociated with Accutase (37° C, 5 min) to yield a single-cell suspension. For each differentiation, ∼730,000 (12-well format) or ∼400,000 (24-well format) hiPSCs were collected and resuspended in the differentiation medium supplemented with ROCKi (10 μM). Cells were evenly spread and maintained at 37 °C with 5 % CO2. Medium was replaced every day with fresh differentiation induction medium. At day six or seven of differentiation, cells were collected for the downstream assays.

For the NGN2-driven directed differentiation, ∼1 x 10^6^ (6-well format) or ∼4 x 10^5^ (12-well format) dissociated single cells were plated on vitronectin coated plates in pre-differentiation medium (Knockout DMEM/F12 (Gibco; 12660012) supplemented with NEAAs, 1x N2 supplement (Gibco; 17502048), 10 ng/ml NT-3 (Peprotech; 450-03), 1 μg/ml laminin, 10 μM ROCKi and 2 μg/ml doxycycline hyclate (Sigma-Aldrich; D9891). The following day, a full medium change was performed in pre-differentiation medium without ROCKi. Two days after doxycycline induction, pre-differentiated neurons were collected as a single-cell suspension via Accutase (37° C, 2 min), counted and plated at the same density onto 6-well or 12-well plates in neural differentiation medium (NDM: half volume DMEM/F12 (Gibco; 11330032), half volume Neurobasal-A (Gibco; 10888022) supplemented with 1x NEAAs, 0.5x GlutaMAX, 1x B-27 minus vitamin A (Gibco; 12587010), 0.5x N2, 10 ng/ml NT-3, 1 μg/ml laminin, and 1 μg/ml doxycycline). Cells were evenly spread, and a full medium change was performed the next day with NDM (without doxycycline). Cells were maintained at 37 °C with 5 % CO2 with half medium changes twice per week until harvest for downstream assays.

### Lentivirus production

Lentivirus production was performed using a second-generation lentiviral system and HEK-293T cells for packaging. Briefly, early passaged HEK-293T cells at 70 – 80 % confluency were transfected with the lentiviral transfer plasmid (37.5 μg per 150 mm flask) as well as with a Pax2 packaging plasmid (Addgene #12260, 20 μg per 150 mm flask) coding for the viral proteins and a MD2.G plasmid (Addgene #12259, 9 μg per 150 mm flask) coding for the VSV-G envelope. To prevent lysosomal degradation and inhibit autophagy, the transfection medium (DMEM, high glucose with 10 % FBS, 2 mM GlutaMax, and 100 units/ml penicillin/streptomycin) was supplemented with 25 μM chloroquine (Sigma-Aldrich; C6628). After 12 – 16 hours, the transfection medium was replaced with the virus collection medium (DMEM, high glucose with 10 % FBS, 2 mM GlutaMax and 100 units/ml penicillin/streptomycin, 1.5 ml of 0.1 N NaOH, and 1 mM sodium butyrate (Sigma Aldrich; 303410)). Viral supernatant was filtered through a 0.45 μm PVDF filter and concentrated via ultracentrifugation using a SW 32 Ti rotor (Beckman Coulter, 107,000x g, 4 °C, 2.5 hours). Viral particles were resuspended in ice-cold PBS or E8 medium with ROCKi (10 μM) (∼500x concentration). All experimental procedures for lentivirus production were performed in a biosafety level 2 laboratory.

### Cell cycle assay

For the quantitative acquisition of 5-ethynyl-2’-deoxyuridine (EdU) incorporation, the ClickTech EdU Cell Proliferation Kit 488 Sensitive (Carl Roth; BCK-EdUPro488FC50) was used. In brief, hiPSCs were incubated for 1 hour in E8 medium supplemented with ROCKi (10 μM) and 10 μM EdU. For NPC labelling, 20 μM Edu (1 hour incubation) was used, while for induced neuron 20 μM EdU was used with extended incubation (2.5 hours) at 37° C. For each staining, 1 x 10^5^ single dissociated cells were used. Cells were fixed in 100 μl of 4 % PFA in PBS and incubated 15 minutes at room temperature in the dark. After quenching with excess PBS with 1 % BSA, cells were centrifuged, and the pellet resuspended in 1X saponin-based permeabilization buffer (200 μl). Staining was performed in 500 μl of the Click reaction solution (RNAse-free water, 1X reaction buffer, catalyst solution, dye azide, and 1X buffer additive) with 30 minutes incubation at room temperature in the dark. Subsequently, cells were washed once with 3 ml of saponin-based permeabilization buffer and stained with DRAQ7 for DNA content (7 minutes incubation at room temperature). Finally, cells were washed with the saponin-based permeabilization buffer and resuspended in 300-500 μl of the saponin-based buffer and EdU incorporation was acquired on the flow cytometer, capturing the fluorescence intensities in the FITC channel (488/20 nm). For the analysis, EdU incorporation was assessed on the logarithmic scale, while the linear scale was used to assess DNA content. Mean fluorescence intensities or percentage of cells in a specific gating group were compared to the experimental control (hiPSCs, NPCs, or induced neurons with NT-ctrl gRNAs).

For the phenotypic EdU-incorporation screen, hiPSCs, NPCs and induced neurons with transduced with the pool of gRNAs were gated for cells in G1-, S-, or G2-phase and sorted >1,000- fold representation. Cells were washed with PBS and pellets snap frozen in liquid nitrogen for subsequent genomic DNA extraction and Amplicon-seq library preparation of gRNAs.

### Reactive oxygen species labelling

To detect the accumulation of ROS in hydrophilic and lipophilic compartments, stainings were performed with 2,7-dichlorodihydrofluoroscein diacetate (DCFA; Cayman Chemical; 601520) and Bodipy C11 (Thermo Fisher Scientific, D38619). The stainings were performed following the guidelines described by Martinez *et al.*^108^. In brief, cells were incubated in a mixture of DCFDA and Bodipy C11 at a final concentration of 5 μM in 1X Hank’s Balanced Salt Solution (Gibco; H9269) for 30 minutes at 37 °C. Afterwards, cells were washed with PBS, dissociated, and resuspended in PBS supplemented with 2 % serum replacement (Gibco; 10828010), 20 μM ROCKi, and DRAQ7 for exclusion of dead cells. ROS content was quantified via flow cytometry (FITC channel – 488 nm), setting gates based on the intensities of unstained cells.

For the phenotypic ROS screen, cells were binned into fluorescently low (0 – 30 % of distribution), medium (35 – 70 %) and high (75 – 100 %) groups. Cells were sorted above a gRNA representation of 1,000x per fraction, washed with PBS and cell pellets snap frozen in liquid nitrogen for subsequent genomic DNA extraction and Amplicon-seq library preparation of gRNAs.

### Amplicon-seq library preparation of the phenotypic CRISPRi screens

To verify the library complexity and to compare gRNA distributions across the phenotypic bulk CRISPRi screens, amplicon-seq libraries were constructed using primer pairs that were targeting flanking sequence of the gRNA cassette in in the modified CROP-seq-opti backbone and that contained illumina adapter sequences required for flow-cell attachment. In addition, to increase sequence diversity, staggered sequences (1 – 6 nucleotides) to the forward (P5) primers were introduced. Libraries were constructed using a mixture of the P5 and sample specific reverse, or P7 primer (primer sequences in **Supplementary Data File 6**). Genomic DNA was extracted using the Quick-DNA Miniprep Plus Kit (Zymo; D4068), following the manufacturer’s instructions and eluting samples in 200 μl of pre-warmed EB buffer to maximize yield. Based on the yield, the library representation was estimated (between 500 – 9000) and multiple PCR reactions for each sample were set up with 0.2 – 1.25 μg input gDNA. The gRNA cassette was amplified from the gDNA using a two-step PCR (initial denaturation (98 °C, 3min) followed by 27 cycles of denaturation (98 °C, 30 sec) and combined annealing/extension (72 °C, 45 sec) using the 2X NEBNext High Fidelity Master Mix (New England Biolabs; M0541). Next, PCR reactions for each sample were pooled and purified with a 1x SPRI magnetic bead cleanup eluting final libraries in EB buffer. Library concentrations were measured using the Qubit (HS DNA; Thermo Fisher Scientific) and the fragment distribution was determined using a high-sensitivity DNA chip on the Bioanalyzer (Agilent) following the manufacturer’s instructions. Finally, libraries were pooled equimolarly and sequenced on the MiSeq or NextSeq 500 platforms with 55 cycles Read 1 and 8 cycles Index 1 (single-end read).

### Analysis of the phenotypic CRISPRi screens

Count matrices of gRNA abundances were constructed by mapping sequencing reads to the *in-silico* reference gRNA library. To compute enrichment scores and significance values, the MaGeCK pipeline was used^102^. In short, the pipeline median-normalizes read counts from each library to account for differences in library size and read count distribution of the multiplexed pool. Next, the mean variance of the read counts was estimated, and a negative binomial model used to test whether there were significant differences in the gRNA abundances in the perturbed samples versus the distribution of read counts in the experimental control. Subsequently, the gRNAs were ranked based on their significance value. According to this rank, the significance testing was collapsed onto the gene-level, computing significance scores for positively and negatively enriched hits. For the phenotypic proliferation screen, the baseline gRNA distribution at day 0 was used as control. For the FACS-based screens, gRNAs derived from unsorted cells was used as control (EdU incorporation screen) or the opposing experimental group was used to compute fold changes in gRNA abundance and significance values (e.g., ROS^high^ versus _ROS_low/mid_)._

### RNA extraction, cDNA synthesis and quantitative real-time PCR (qPCR)

Cells were harvested washing them twice with PBS and dissociating them with Accutase (37 °C, 3 – 5 min incubation). The cell suspension was collected in PBS and pellets snap-frozen in liquid nitrogen. An RNA extraction kit (RNAeasy Mini, Qiagen; 74106) was used in combination with a cell-lysate homogenizer (QIAshredder, Qiagen; 79656) following the manufacturer’s instructions. To eliminate residual genomic DNA, on-column DNase treatment was applied (Qiagen; 79254). Subsequently, cDNA was generated using the High-Capacity cDNA Reverse Transcription Kit (Thermo Fisher Scientific; 4368814) following the manufacturer’s instructions.

Quantitative real-time PCR was performed using the Power SYBR® Green PCR Master Mix (Thermo Fisher Scientific; 4367659) on a StepOnePlus Real-Time PCR system (Applied Biosystems). Reactions were set in technical triplicates with qPCR primers that span exon-exon junctions (sequences obtained from PrimerBank^109^, listed in **Supplementary Data File 6**). Cycle threshold (Ct) values were computed with the StepOne software (v2.3), reviewed for successful PCR reactions and averaged. Relative gene expression was quantified according to the ΔΔCt method in respect of two housekeeping genes (*GAPDH*, *ACTB*) and the experimental control sample.

### CRISPRi screen coupled to scifi-RNA- and scifi-ATAC-seq

#### Virus transduction, cell preparation and nuclei isolation

4.5 x 10^6^ monoclonal CRISPRi-NGN2-hiPSCs were transduced with the gRNA pool (*n* = 275) targeting 65 schizophrenia implicated genes at an MOI of ∼0.05 (verified via flow cytometry based on tagBFP expression) at a minimum representation of ∼1,000 cells per gRNA. Cells were selected via puromycin treatment 24 hours post-transduction for three days (day 1: 1 μg/ml, day 2: 2.5 μg/ml, day 3: 3 μg/ml). Selected cells were recovered for 24 hours and the pool of cells split into three groups at a representation >1,000 cells per gRNA per group. One group of cells was maintained at the hiPSC stage, while the other two were differentiated to either NPCs or induced neurons as described above. At day 11 post-transduction, NPCs and at day 12 induced neurons were collected and processed with the scifi-RNA- and scifi-ATAC-seq workflows. Nuclei were isolated as described above and fixed in a 3 % glyoxal solution (40 % glyoxal (Merck; 128465), 20 % ethanol, 0.75 % acetic acid; adjusted to pH 4 by addition of 1 M NaOH), incubating the sample for 7 minutes at room temperature. Afterwards, fixation was stopped by adding excess of PBS with 2 % BSA, nuclei centrifuged (300 x g, 5 min) and washed twice with PBS with 2 % BSA before proceeding with pre-indexing.

#### High-throughput scRNA-seq library preparation with scifi-RNA-seq

The scifi-RNA-seq workflow was followed as previously described by Datlinger *et al*.^46^. Briefly, nuclei were counted and split into 24 bulk pools of ∼20,000 nuclei each. Pre-indexing barcodes were introduced via *in-situ* reverse transcription mRNA via barcoded poly-dT primer (**Supplementary Data File 6**) using 50 units Maxima H Minus Reverse Transcriptase (Thermo Fisher Scientific; EP0753) on a thermocycler with the following program: 50 °C for 10 min, followed by 3 thermal cycles (8 °C for 12 sec, 15 °C for 45 sec, 20 °C for 45 sec, 30 °C for 30 sec, 42 °C for 2 min and 50 °C for 3 min), and final incubation at 50 °C for 5 min. The reverse transcription reaction was spiked-in with gRNA-specific primers that included matching pre-index sequences and were targeting the gRNA scaffold (**Supplementary Data File 6**) to increase gRNA recovery. Afterwards, all nuclei were pooled, washed in PBS with 2 % BSA, strained through a 40 μm mesh filter (Falcon) and additionally through a FlowMi filter and counted via DAPI staining (Countess 3 FL). The thermoligation reaction was prepared (1X ampligase buffer, ampligase (230 units; Lucigen; 112501), Reducing Agent B, and 230 pmol/l bridge oligonucleotide (5’- CGTCGTGTAGGGAAAGAGTGTGACGCTGCCGACGA[ddc]-3’)) and added to the nuclei (in 1X ampligase buffer). For scifi-RNA-seq, ∼160,000 and ∼250,000 matched nuclei from NPCs and neurons were collected and nuclei were subjected to the 10x Chromium controller for single nucleus partitioning as described above (Chip H, 40 μl partitioning oil dispensed on row 3 instead of 45 μl as described above for Chip J) and the cell barcoding reaction performed on a thermocycler with the following program (lid temperature 98 °C): 12 cycles of denaturation at 98 °C for 30 sec followed by annealing and ligation at 59 °C for 2 min. GEMs were broken and the sample purified as described above using Dynabead MyOne SILANE beads and subsequently SPRI beads. Next, template switching was performed incubating the purified cDNA in template switch reaction solution (template-switch oligo (2.5 μM; 5’- AAGCAGTGGTATCAACGCAGAGTGAATrGrGrG-3’), 1X reverse transcription buffer (Thermo Fisher Scientific), dNTPs (50 μM each), 4 % FicoII PM-400 (Sigma-Aldrich; F4375), Protector RNAse-inhibitor (50 units, Roche; 3335402001), and Maxima H Minus Reverse Transcriptase (500 units, Thermo Fisher Scientific)) on a thermocycler with an initial incubation at 25 °C for 30 min followed by an incubation at 42 °C for 90 min. Template-switched cDNA was purified with SPRI beads (1.2X ratio) and full-length cDNA was amplified using a universal partial P5 primer (500 nM), the TSO enrichment primer (500 nM; 5’-AAGCAGTGGTATCAACGCAGAGT-3’) and the NEBNext High-Fidelity 2X PCR master mix (New England Biolabs) with the following program: initial denaturation (98 °C, 3 min) followed by 5 cycles of 98 °C, 30 sec (denaturation), 65 °C, 30 sec (annealing), and 72 °C, 3 min (elongation). After the five initial cycles, a fraction of amplified material was quantified using SYBR Gold and qPCR (StepOne plus) in order to determine the additional number of amplification cycles. A final elongation step at 72 °C for 5 min was added and samples were purified using SPRI beads (0.8X clean-up followed by an additional 0.6X clean- up). Full-length cDNA was quantified using the Qubit HS-DNA assay (Thermo Fisher Scientific) and the quality evaluated on the Bioanalyzer (HS-DNA kit, Agilent).

Approximately 25 % of full-length cDNA was used to construct the final whole-transcriptome scRNA-seq library, following a Tn5-based tagmentation approach. In short, the Tn5 molecule was prepared by symmetrical loading with oligonucleotides carrying a universal primer binding site (Nextera Read 2) that allows the subsequent amplification and construction of illumina-compatible libraries. For Read2N-trasnposome assembly, 100 μM of the mosaic end-complement oligonucleotide (ME-ddc, 5’-[Phos]CTGTCTCTTATACACATCT[ddc]-3’) was mixed with 100 μM of a universal Nextera Read 2 oligonucleotide (5’- GTCTCGTGGGCTCGGAGATGTGTATAAGAGACAG-3’) and annealed in 1X annealing buffer (10 mM Tris-Hcl (pH 7.5), 50 mM Nacl, and 1 mM EDTA) by incubating for 95 °C for 3 minutes followed by cooling down to 25 °C at a ramp rate of -1 °C/min. Annealed oligonucleotides were diluted in an equal volume of glycerol (Merck). Transposomes were assembled mixing equal volumes of the annealed and diluted oligonucleotides with in-house produced Tn5 molecules (diluted to 1 mg/ml in 50 mM Tris, 100 mM NaCl, 0.1 mM EDTA, 1 mM DTT, 0.1 % NP-40, and 50 % glycerol) incubating at room temperature for 30 – 60 min. Transposition was performed with 1 – 10 ng amplified full-length cDNA in 1X reaction buffer (10 mM TAPS, 5 mM MgCl2; adjusted to pH 8.5 with NaOH and ∼0.94 μM Tn5) at 55 °C for 10 min. The transposition reaction was quenched with 0.1 % SDS and 5 min incubation at room temperature. An equal volume of water was added to the sample and tagmented cDNA was purified via 1X SPRI clean-up. Final scifi-RNA-seq libraries were prepared using universal P5 and indexed P7 primer pairs (500 nM each) with the NEBNext High-Fidelity 2X PCR master mix (New England Biolabs) on the thermocycler with the following amplification program: initial denaturation at 98 °C, for 30 sec followed by 12 cycles of 98 °C, for 10 sec (denaturation), 65 °C, for 30 sec (annealing), and 72 °C for 1 min (elongation). The PCR product was purified using a 0.7X ratio of SPRI beads. Libraries were quantified and the quality evaluated via Qubit and Bioanalyzer as described above. The scifi-ATAC-seq libraries were sequenced on the NovaSeq6000 (v1.0) platform (Read 1: 19 cycles, Read 2: 90 cycles, Index 1: 8 cycles, Index 2: 16 cycles).

Another 25 % of full-length cDNA was used to construct the targeted-sequencing library using a hemi-nested PCR approach as described previously by Schraivogel *et al*.^44^ (inner and outer primer pool sequences are listed in **Supplementary Data File 6**). For the design of the pool, the R package TAPseq (v1.14.1) was used. The first two rounds of hemi-nested PCR were performed using the oligonucleotide pools (1.5 μM final) with a universal P5 primer (partial P5, 500 nM) and the NEBNext 2X High-Fidelity PCR master mix (New England Biolabs) and following the thermal amplification program (lid temperature 98 °C): 98 °C, for 30 sec followed by 8 cycles of 98 °C, for 10 sec (denaturation), 69 °C, for 30 sec (annealing), and 72 °C for 30 sec (elongation), and final extension at 72 °C, for 2 min. The final PCR reaction was performed using equimolar amounts of the universal P5 primer (partial P5, 500 nM) and library indexed TruSeq P7 primer (500 nM) and the standard final PCR reaction as described for the whole-transcriptome scifi-RNA-seq libraries (annealing at 65 °C, 30 sec and elongation at 72 °C for 1 min). Libraries were purified and the quality as described via Qubit and Bioanalyzer. The TAP-seq libraries were sequenced on the NextSeq2000 platform (Read 1: 19 cycles, Read 2: 90 cycles, Index 1: 8 cycles, Index 2: 16 cycles (custom sequencing primer used for Index 2 read: 5’-GGGAAAGAGTGTGACGCTGCCGACGA- 3’)).

Another ∼20 % of the full-length cDNA was used to construct the gRNA enrichment libraries adopting the hemi-nested PCR approach from Hill *et al.*^43^ to the scifi-RNA-seq library structure. Briefly, gRNA transcripts were amplified from full-length cDNA using a universal P5 primer and in three consecutive rounds of PCR with primers annealing to constant hU6 promoter sequences of gRNA transcripts (primer sequences listed in **Supplementary Data File 6**) using the KAPA HiFi HotStart mix (Roche, KK2602). The first two hemi-nested PCRs were performed at 65 °C annealing for 10 s followed by extension at 72 °C for 15 s. The last PCR was performed combining annealing and extension steps at 72 °C for 45 s. Guide RNA enrichment libraries were sequenced on a MiSeq using standard illumina primer (Read 1: 19 cycles, Read 2: 43 cycles, Index 1: 8 cycles, Index 2: 16 cycles).

#### High-throughput scATAC-seq library preparation with scifi-ATAC-seq

The scifi-RNA-seq workflow was modified following the same basic principle of combinatorial fluidic indexing and adjusted to profile chromatin accessibility with high-throughput at the single- cell level. Briefly, transposomes for *in-situ* tagmentation of open chromatin regions were assembled by preparing annealed oligonucleotides for loading. In short, 100 μM of universal Nextera Read 1 oligonucleotides (5’-TCGTCGGCAGCGTCAGATGTGTATAAGAGACAG-3’) and 100 μM of oligonucleotides with a Nextera_Read2-sample_pre-index structure (5’- CAAGCAGAAGACGGCATACGAGAT[pre-index] GTCTCGTGGGCTCGGAGATGTGTATAAGAGACAG-3’; pre-indices listed in **Supplementary Data File 6**) were annealed with 100 μM of the mosaic end-complement oligonucleotide at a 1:1:2 ratio by heating for 95 °C for 3 minutes and cooling down to 25 °C at a ramp rate of -1 °C/min in 1X annealing buffer (10 mM Tris-Hcl (pH 7.5), 50 mM Nacl, and 1 mM EDTA). Annealed oligos were mixed with an equal volume of glycerol (Merck) and stored at -20 °C until use. Transposomes were assembled using the in-house produced Tn5 (diluted to 1 mg/ml in 50 mM Tris, 100 mM NaCl, 0.1 mM EDTA, 1 mM DTT, 0.1 % NP-40, and 50 % glycerol) and the prepared, annealed and diluted via incubation at room temperature for 30 min.

During transposome assembly, nuclei, derived from HEK-293T, NIH-3T3, NPCs or induced neurons were isolated as described, fixed (3 % glyoxal incubation at room temperature followed by washes in PBS with 2 % BSA) and counted via DAPI staining (Countess 3 FL). Afterwards, the pre-index was introduced to open chromatin regions via *in-situ* tagmentation of split bulk pools of nuclei (∼20,000) in a transposition reaction solution containing 38.8 mM Tris-acetate, 77.6 mM potassium acetate, 11.8 mM magnesium acetate, and 18.8 % dimethylformamide supplemented with 0.005x protease inhibitor cocktail (Roche), 0.4 U/μl SUPERaseIn (Thermo Fisher Scientific), 1.2 U/μl RiboLock (Thermo Fisher Scientific) and the Tn5 transposomes with the pre-indexing barcodes (1:10 (v/v); 938 nM) and incubated at 30 °C for 37 min with shaking at 400 rpm. Split pools were washed with PBS with 2 % BSA. To recover gRNA identities from the scATAC-seq readout, a reverse transcription was performed using primers with matching pre-indexing barcodes (500 nM final) in a RT reaction mix with 1 mM dNTPs, 20 units Protector RNase-inhibitor (Roche) and 200 units RevertAid reverse transcriptase (Thermo Fisher Scientific; EP0441 ) with 30 min incubation at 37 °C. Afterwards, nuclei were pooled, washed with excess PBS with 2 % BSA, filtered and counted as described for the scifi-RNA-seq workflow. Approximately 90,000 and ∼140,000 nuclei were mixed with the cell barcoding reaction solution (Barcoding Reagent B, Reducing Agent B, Barcoding enzyme and 1.2 μl of a 50 μM spike-in primer to exponentially amplify gRNA-derived cDNA) using the Chromium Next GEM Single Cell ATAC Library & Gel Bead Kit (v1) (10x Genomics; 1000176) and the 10x Chromium controller was loaded as described above. After single nucleus partitioning, the sample was transferred onto a thermocycler for the barcoding reaction (72 °C, for 5 min (gap repair); 98 °C, for 30 sec (initial denaturation) followed by twelve cycles of 98 °C, 10 sec (denaturation), 59 °C, for 30 sec (annealing), and 72 °C, for 1 min (elongation). Subsequently, GEMs were broken and purified as described above. Final scifi-ATAC-seq libraries were constructed using universal P5/P7 primers (500 nM each) and the NEBNext High-Fidelity 2X master mix (New England Biolabs) with the following PCR program: 98 °C, 30 sec (initial denaturation), followed by six cycles of 98 °C, 10 sec (denaturation), 65 °C, 30 sec (annealing), and 72 °C, for 1 min (elongation). Libraries were purified using 1.2X SPRI magnetic beads. Quantity and quality of the libraries were assessed via Qubit and Bioanalyzer as described above. Libraries were sequenced on the NovaSeq6000 platform (Read 1: 50 cycles, Read 2: 50 cycles, Index 1: 11 cycles, Index 2: 16 cycles). From a fraction of the final scifi-ATAC-seq library, gRNA enrichment libraries were constructed using a modified enrichment protocol to the one described above. Oligonucleotides for the nested PCR were biotinylated and gRNA enriched fragments were purified using Dynabeads MyOne Streptavidin T1 (Thermo Fisher Scientific; 65601) following the instructions of the manufacturer. Enrichment libraries were sequenced as described above.

### Pre-processing and downstream analysis of scifi-RNA-seq

Sequencing reads were demultiplexed based on the pre-index using Je (v2.0.RC), while permitting two mismatches. The 10x cell barcode (Index 2 read) was appended to Read 1 and sequencing reads mapped to the reference genome (hg38) using STARsolo (v2.7.9) with the settings that most resemble CellRanger. For each cell type cell calling was performed by determining inflection points on a rank vs UMI or gene plot (>= 600 UMI and 500 genes/cell for NPCs and >= 700 UMI and 600 genes/cell for neurons). Additionally, cells were filtered by mitochondrial read percentage (<=5 %) and ribosomal read percentage (<= 10 %). Gene expression counts were normalized with SCTransform.

Guide RNA libraries were demultiplexed based on the pre-index using Je and mapped to the in-silico library. Next gRNA identities were assigned to individual cells using both fba (v0.0.11) and kite workflow based on bustools (v0.41.0) and kallisto (v0.46.2). We assigned a gRNA to a cell if there was at least one gRNA read in either method.

To evaluate the overall effect of CRISPRi perturbations on the transcriptomic readouts, the R package scPerturb was used (v0.1.0). For this, the distance ("E-distance") between a group of cells assigned to a particular perturbation was compared to a group of control cells in the latent space and a score was computed. Next, the significance of a given E-distance was tested by calculating a p-value and comparing it to 1000 empirical permutations of random cells of the same size. The E-distances between cells with a particular gRNA and all non-targeting gRNAs were calculated in the principal component space of the RNA experiment in NPCs and induced neurons separately. Guide RNA identities were filtered to be present in at least five cells in each respective cell type. A lenient significance threshold was applied at the gRNA level (E-distance *P* < 0.3) and retained gRNAs were collapsed onto the gene level by calculating the E-distances between cells with gRNAs for the same gene and the non-targeting gRNAs. For **Fig. 1d**, external datasets^14,15,44,96–101^ from the resource were used to compare the effect sizes of our perturbations.

**Fig. 1e** shows the estimates of the E-distance between the perturbation and non-targeting control cells and are colored by the E-distance significance (FDR < 0.1).

To identify gRNAs that were enriched in each cluster, we performed fisher tests comparing the prevalence of guides (collapsed per gene target) in each cluster to the other clusters in that cell type (*P* ≤ 0.05).

Differentially expressed genes for each cluster were identified by contrasting the transcriptional profiles with all other cells from the same cell type (NPC or neuron) and using the Wilcoxon test adaptation in Seurat with *FindMarkers* (average log2FoldChange threshold ≥ ±0.2, transcripts expressed in at least ten cells).

To increase power of differential gene expression analysis, gene set modules were constructed based on gene expression similarities across CRISPRi perturbation using Monocle 3^58^ with *find_gene_modules* using standard parameters.

### Pre-processing of scifi-ATAC-seq

Sequencing reads were demultiplexed based on the pre-index using fastq-multx, while permitting two mismatches. Demultiplexed reads were aligned to the hg38 genome and fragment files were generated using chromap (v0.2.3). For the mixed-species benchmarking experiment, a combined hg38/mm10 reference genome was used. Fragments mapping to either of the genomes were counted in order to assess collision rates. Using the R package ArchR (v1.02), fragment counts were computed for each cell in 500 base-pair bins to generate a celll-by-bin matrix.

Cell calling was performed for each cell type according to the number of unique fragments per cell (≥ 2000 fragments per cell), TSS enrichment score (≥ 3), and fraction of reads in promoter regions (≥ 0.02). Peak calling was performed for each cell type separately on clusters called by louvain clustering on the cell-by-bin matrix. Shortly, the MACS2 peak caller (v2.2.7.1) was used to identify fixed-with 501-bp peaks (params; *–call-summits --keep-dup all –shift -75 –extsize 150 –q 0.05*) for each cluster of cells. Afterwards, an iterative overlap peak merging procedure was performed to derive a consensus peak set for all samples^110^. From here a peak by cell matrix was counted and normalized by TF-IDF normalization.

To infer TF motif activity we first linked TFs to ATAC peaks by performing *in silico* ChIP-seq^111^. In brief, for each ATAC peak containing a TF motif we correlate peak accessibility with corresponding TF RNA expression. This correlation is then weighted for motif score for that peak as well as the peak maximum accessibility across the dataset, and finally minmax normalized. This value is then used to filter TF-motif/peak links (cutoff = 0.05). To allow for correlating the single-cell readouts we infer metacells from the ATAC readout with SEAcells^112^ and transfer the metacell annotation to the RNA readout, resulting in peak- and gene-by-common metacell matrices. The derived motif matches were then used as input for chromVAR to calculate motif activity scores (deviations and z-scores)^113^.

### Integration and co-embedding of single-nucleus chromatin accessibility and gene expression readouts

To integrate single-cell chromatin accessibility and gene expression data, a graph-based neural network approach (GLUE) was used to infer a shared embedding (50 latent variables). To model cross-modality relations despite differing semantics, GLUE utilizes a user-provided graph representation of links between ATAC- and RNA-seq data. Here, the graph contains connections between gene expression (RNA) and respective gene body accessibility peaks (ATAC).

Additionally, 1) a gene-by-cell matrix of highly variable genes and a 100 PC representation of the gene expression data is provided as well as 2) a peak-by-cell matrix of peaks connected to highly variable genes in the graph and a 100 LSI representation of the peak data. Adversarial multimodal alignment of the cells is performed as an iterative optimization procedure, guided by feature embeddings encoded from the user-provided graph. Hyperparameter optimization was done by evaluating integration consistency - a set of mean correlations for graph connections (ATAC peak

∼ gene expression), weighted by the graph edge weight, for various size groupings of cells (k-means clusters of 10, 20, 50, 100, and 200 cells). A higher curve (on a metacell vs consistency score plot) indicates better integration, with minimum threshold at 0.05 as suggested by the developers of GLUE.

The inferred latent variables were used to compute a neighborhood graph, which was further used to compute a UMAP and perform leiden clustering. Clustering resolution was determined by evaluating intercluster homogeneity and cluster to cluster separation, adapting an approach from Persad *et al.* 2024^112^. Briefly diffusion components were computed using the latent variables, whereafter per cluster variance in diffusion components was computed to determine intercluster homogeneity (lower values indicate high homogeneity of a cluster), and diffusion distance between nearest neighbord clusters for each cluster to define a value for cluster separation (high values indicate higher distinction between clusters).

### Multiome timecourse data generation and processing from parental hiPSCs, NPCs, and induced neurons over the course of differentiation

Human iPSCs expressing the CRISPRi machinery (*n =* 2 biological replicates) were either differentiated to NPCs or induced neurons as described above. NPCs were collected seven days after differentiation induction and induced neurons over a timecourse of 1, 2, 5, 7, 14, 21, and 28 days via Accutase (7 min (NPC) or 3 min (induced neurons) at 37° C), washed twice with PBS and resuspended in 1X lysis buffer (10 mM Tris-HCl pH 7.5, 10 mM NaCl, 3 mM MgCl2, 0.1 % Tween20, 20 μg/ml RNase-in, 1 mM DTT (Roche; 10708984001), 1 % BSA, 0.025 % IGEPAL CA-630 and 0.001 % Digitonin (Thermo Fisher Scientific; BN2006) (NPCs) or 0.01 % Digitonin (induced neurons)) and incubated for 5 minutes on ice. The reaction was quenched with wash buffer (10 mM Tris-HCl pH 7.5, 10 mM NaCl, 3 mM MgCl2, 0.1 % Tween20, 20 μg/ml RNase-in, 1 mM DTT, 1 % BSA), nuclei resuspended in PBS with 1 % BSA and filtered through a 40 μm mesh filter (Falcon) and counted via DAPI staining on a Countess 3 FL. From this point, library preparation was performed according to the manufacturer’s instructions (Chromium Single cell Multiome ATAC + Gene Expression; 10000285). Briefly, nuclei were resuspended 1X Nuclei Buffer (provided with the kit) and the tagmentation reaction mix was added to the nuclei suspension. *In-situ* tagmentation was performed for 60 min at 37 °C on a thermocycler (lid temperature: 50° c). Immediately after tagmentation, the barcoding reaction mix was added to the nuclei and the sample was loaded on the Chromium controller (10x Genomics) for single nucleus partitioning as described by the manufacturer. In brief, empty lanes on the chip (Chip J) were filled with 50 % glycerol (row 1: 70 μl, row 2: 50 μl, row 3: 45 μl) and the sample was pipetted slowly onto row 1 (70 μl). Gel beads were vortexed for 30 sec, briefly centrifuged, checked for potential bubbles and added to row 2. Finally, 45 μl of partitioning oil was added to row 3. Next, cell barcoding was performed on the thermocycler with the following program (lid temperature: 50 °C): 37 °C for 45 min, 25 °C for 30 min and hold at 4 °C. GEMs were broken with the addition of 125 μl Recovery Agent and the soluble phase was purified via Dynabeads (MyOne SILANE). After an additional 1.8X SPRI cleanup, libraries were pre-amplified and physically split in two for the construction of modality-specific libraries. Final libraries were quantified via Qubit (HS-DNA) and the fragment distribution assessed via a Bioanalyzer run (HS-DNA). Single-cell ATAC- and scRNA-seq libraries were sequenced on the NextSeq2000 platform with the following read configurations: ATAC Read 1: 50 cycles, ATAC Read 2: 49 cycles, ATAC Index 1: 8 cycles, ATAC Index 2: 16 cycles; RNA Read 1: 28 cycles, RNA Read 2: 90 cycles, RNA Index 1: 10 cycles, RNA Index 2: 10 cycles.

The multiome dataset was pre-processed with STARsolo (RNA modality) and chromap (ATAC modality) mapping the respective reads to the human reference genome (hg38). Gene expression count matrices were normalized using Seurat (SCTransform) and cell calling applied using a threshold of minimum 500 genes per cell and maximum 11,500 (to remove doublets), ≤ 15 % mitochondrial and ≤ 15 % ribosomal reads for the scRNA modality. For the scATAC readout, a cell matrix was generated using ArchR. Cell calling thresholds were applied (> 1,000 fragments per cell, > 0.12 fraction of peaks in promoter regions, and TSS score > 3). Peak counts were normalized using TF-IDF normalization followed by dimensionality reduction with iterative LSI and SVD. A 50 shared latent variable representation was constructed with GLUE as described in the previous section and used as input to construct a partition-based graph abstraction (PAGA) using *sc.tl.paga()*, with clusters defined by the Leiden algorithm. The layout was intialized for visualization by recalculating the graph layout with *sc.tl.draw_graph(init_pos=’paga’)*. To perform diffusion pseudotime inference, we specify a root cell in the iPSC-like Leiden cluster using iroot, which defines the starting point for pseudotime calculation. Finally, we calculate pseudotime with *sc.tl.dpt()*. To assign cells to nodes along the pseudotime trajectory, we constructed a principal graph with ElPiGraph (nodes=15) using latent variables and the PAGA graph as input.

### Enrichment analysis and comparisons with transcriptional signatures

Protein-protein interactions were derived from STRINGdb (webtool, v12.0). We downloaded differentially expressed genes in neuronal subsets from post-mortem brains of schizophrenia patients versus controls from Ruzicka *et al.*^18^ . Genes that attained adjusted p-value < 0.05 were used as input for scDRS^105^ to compute significance per cell and enrichment at the cluster level, using the logFC as weights.

Transcriptional signature scores were computed using the *AddModuleScore* function of the R package Seurat and averaged the mean expression of associated genes for each cluster. The cell cycle markers were derived from Kowalczyk *et al*^70^. The glycolysis signature score was computed according to genes listed in the gene ontology database (biological process; GO:0006096).

Overrepresentation analysis was performed with clusterProfiler (v4.0.2) using terms retrieved from the Gene Ontology, the REACTOME, the KEGG, the DisGeNET, and the Molecular Signatures databases (MSigDB). For enrichment of GWAS-derived traits, the tool FUMA was used (1.5.2) with the function “GENE2FUNC”, and correcting raw *P* values according to Benjamini-Hochberg (FDR cut-off ≤ 0.05). TF associations were retrieved using ChEA3^103^.

### Gene regulatory network inference

Transcriptome-based gene regulatory network analysis was performed using SCENIC^73^. Briefly, the expression matrix was filtered to retain genes that are expressed in at least 1 % of the cells, log-normalized and a co-expression matrix was constructed running GENIE3 (SCENIC::*runGenie3*). Regulons were retrieved from the RcisTarget package. Motif enrichment was performed 10 kilobases up- and downstream of gene promoters. TF-gene connections were ranked according to weight and hub TFs identified after filtering the dataset for the top 5000 connections.

GRN inference using single-cell gene expression and chromatin accessibility readouts was performed using Single-Cell Deep multi-Omic Regulatory Inference (scDoRI). scDoRI is an end-to-end approach that integrates single-cell RNA and ATAC data using an autoencoder (AE) framework. Drawing from dimensionality reduction and regulatory network inference principles, scDoRI decomposes each cell’s expression and accessibility profiles into a shared set of regulatory “topics,” akin to topic modeling.

A key innovation of scDoRI is the augmentation of the AE’s primary decoding objective with linear auxiliary decoding tasks, guiding inferred topics to capture established enhancer-mediated gene regulatory network (eGRN) signatures^114^. Each topic aligns with active transcription factors (TFs), co-accessible cis-regulatory elements (CREs), and enhancer-gene linkages. When integrated with *in silico* TF binding predictions, these components yield topic-specific eGRNs.

Three core linear decoders underpin this framework: (i) a topics-to-peaks decoder reflecting co-accessibility, (ii) a peak-to-gene matrix identifying putative enhancer-target gene links, and (iii) a topics-by-TF matrix highlighting TF co-expression. By applying a softmax transformation to the latent space, scDoRI also naturally regularizes the number of active topics, enhancing interpretability and enabling the dissection of regulatory networks across continuous cell states.

### Zebrafish mutagenesis experiments

Wild-type AB/AB2B2 zebrafish were used in this study. Animals were housed in accordance with the European Union animal welfare standards and all animal experiments were performed in accordance with the European Union Animal Welfare Guidelines and the EMBL Institutional Animal Care and Use Committee (IACUC code: 22-003_HD_MD). Kimmel^115^ staging was used to stage zebrafish embryos and larvae. Adult fish were fed daily with hatched Artemia and maintained at 28.5 °C with light/dark cycles of 14 to 10 hours. 48 hpf and 72 hpf larvae were used in this study.

Gene sequences were selected using the Ensembl database and gRNAs designed with CRISPOR^116^ (sequences listed in **Supplementary Data File 1**). The gRNAs with the highest efficiency and the fewest off-targets were selected and ordered as duplexes of crRNA/tracrRNAs (Sigma Aldrich). Injections into one-cell stage embryos were performed following Hoshijima *et al*.^117^ with assembled ribonucleoprotein complexes (RNPs) including 5 μM of combined gRNAs and 5 μM of Cas9 protein (produced by the Protein Expression and Purification facility at EMBL). Wild-type AB/AB2B2 and gRNA control (targeting *slc45a2*, derived from Hoshijima *et al*.^117^) animals were used as controls. Embryos were grown in petri dishes with 1x E3 medium (for 60x stock solution: 17.2 g NaCl, 0.76 g KCl, 2.9 g CaCl2 x 2 H2O, 4.9 g MgSO4 x 7 H2O, 1 L distilled H2O) at 28.5°C. At 6 hpf, the embryos were cleared of dead and unfertilized embryos, fresh E3 and methylene blue were added. Fish were reared at 28°C until 72 hpf. At 24 hpf, pronease was added to dechorionate them.

Zebrafish embryos at 48 hpf and 72 hpf were anaesthetized prior to imaging with MS222. A Leica MSV269 stereomicroscope connected to a K3M camera was used for imaging. Images were taken at 2.5x and 5x magnification. Individual anaesthetized fish were embedded in 1.5 % agarose scaffolds and covered with 0.6 % low melting agarose. Head sizes of the animals were measured using Fiji.

For genotyping, we followed Saunders *et al.*^71^. Briefly, three batches of 5x 24 hpf F0-injected individuals were collected. Genomic DNA was extracted from these fish using 100 µl alkaline lysis buffer (25 mmol NaOH, 0.2 mmol EDTA) and incubated at 95°C for 30 min. The same volume of neutralization buffer (40 mmol Tris-HCl, pH 5.0) was then added. The purified PCR product was submitted for Sanger sequencing to assess cutting efficiencies.

### Bulk ATAC-seq

Bulk profiling of chromatin accessibility was performed similarly to its single-cell counterpart with few modifications. Briefly, hiPSCs, and derived NPCs or induced neurons were washed twice with PBS and dissociated via Accutase (3 – 7 min depending on cell type, 37 °C). Cells were counted and 50,000 – 100,000 per tagmentation reaction were resuspended in 1X lysis buffer and incubated for 5 min on ice. Lysis was stopped adding 1 ml of wash buffer as described above and nuclei were centrifuged (500x g, 5 min). The supernatant was discarded, and nuclei resuspended in the Tn5 reaction solution (38.8 mM Tris-acetate, 77.6 mM potassium acetate, 11.8 mM magnesium acetate, and 18.8 % dimethylformamide supplemented with 0.005x protease inhibitor cocktail (Roche), and the Tn5 transposomes with the pre-indexing barcodes (1:10 (v/v); 938 nM)) and incubated at 37 °C for 30 min with shaking at 500 rpm. ATAC fragments were purified using the MinElute DNA purification kit according to the manufacturer’s instructions eluting the DNA in 22 μl pre-warmed EB buffer (70 °C). Finally, illumina-compatible sequencing libraries were constructed using a universal P7 primer (partial P7) and indexed P5 primer and the NEBNext High-Fidelity PCR master mix with the following amplification program: 72 °C for 5 min (gap repair) and 98 °C for 30 sec (initial denaturation) followed by six cycles of 98 °C for 10 sec (denaturation), 65 ° C for 30 sec (annealing), and 72 °C for 1 min (elongation). Bulk ATAC libraries were purified via 1.2X SPRI magnetic beads, quantified via Qubit (HS-DNA kit) and the fragment distribution evaluated via the Bioanalyzer (HS-DNA kit). Libraries were pooled equimolarly, and the multiplexed pool was sequenced on the NovaSeq6000 platform with standard illumina sequencing primer (Read 1: 50 cycles, Read 2: 50 cycles, Index 1: 11 cycles, Index 2: 6 cycles).

### Analysis of the bulk ATAC-seq readouts

Briefly, demultiplexed ATAC-seq reads were mapped to the human genome (hg38) using bowtie2 (v2.3.4.1). Next, peaks were called with MACS2 (v2.2.7.1) with the following parameters: *-- nomodel --shift -75 --extsize 150 -B --SPMR --keep-dup all --call- summits*. Then, DiffBind was used to jointly analyze called peaks and to identify differentially accessible peaks. Peaks were considered for downstream analysis that were detected in at least two samples. A consensus peak set was generated using an overlap of 66 %. A LOESS normalization was applied to correct for counts per peak in all ATAC-seq datasets. Normalized ATAC-seq counts were obtained using edgeR (v3.18) with the following model: *y* ∼ *Treatment*, where Treatment corresponds to the modality of the CRISPR perturbation (activation) and target (*TCF4* or *ZEB1*). Guide RNA #2 targeting *TCF4* was excluded from analysis due to low overexpression capacity. Consensus peak- sets of control samples were used as a reference for comparison. Log2FoldChanges between perturbations and controls were assessed using the exact test (two-sided), differentially accessible regions (DARs) were considered as such when the adjusted p-value < 0.1.

Genes in the proximity of DARs (±50 kilobases) were used to establish peak-to-gene connections. The list of DAR-derived differentially regulated genes was used for the overrepresentation analysis. This was performed with terms retrieved from the Gene Ontology, the REACTOME, the KEGG, the DisGeNET, and the Molecular Signatures databases (MSigDB) using clusterProfiler (v4.0.2).

The DAR-derived gene list was used as input to compute TF activity scores in the scCRISPRi transcriptomic readout (scifi-RNA-seq). For this, the *AddModuleScore* function of the R package Seurat was utilized that calculates the mean expression of the provided genes across all comparison groups (clusters). This average value is normalized according to the aggregated expression of genes that are used as background control.

For the comparison of putative target genes of TCF4, ZEB1, and MCRS1, the Chip-Atlas database was used. Specifically, the following datasets were retrieved: i) for TCF4: SRX1491250 (CAL-1), SRX220960 (CAL-1), SRX1491252 (GEN2.2), SRX702131 (hESC derived ectodermal cells), SRX702132 (hESC HUES64), SRX11919215 (iPS cells), SRX11919216 (iPS cells), SRX2661484 (SH-SY5Y), SRX2661486 (SH-SY5Y), SRX11161903 (SK-N-SH), SRX11161904 (SK-N-SH), SRX2430641 (SW1783); ii) for ZEB1: SRX5457540 (Bipolar spindle neurons), SRX17812782 (Dorsolateral prefrontal cortex), SRX17812784 (Dorsolateral prefrontal cortex); iii) for MCRS1: SRX2708818 (HepG2), SRX2708819 (HepG2), SRX3445802 (HepG2), SRX3445803 (HepG2), SRX2708820 (HuH-7), SRX2708821 (HuH-7).

### Software

Schematics and illustrations were generated with Biorender. Data plots were generated using R and Python and related packages or with Graphpad prism (version 10). Figure panels were assembled with Adobe Illustrator.

## Acknowledgements

We thank Charles Girardot and the Genome Biology Computational Support for their assistance in data analysis and submission. We thank the staff of the European Molecular Biology Laboratory (EMBL) Genomics Core Facility, Protein Expression and Purification Facility, and Flow Cytometry Core Facility for providing essential chemistry, sample handling and data generation. We thank all members of the EMBL facilities who provide essential consumables for experimental laboratory work. We thank Nadine Fernandez Novel-Marx, Sara Lobato Moreno, Tekin Can Sobaci, Minyoung Kim and Shalmalee Kharkar, for maintenance of the hiPSCs lines and helping with sgRNA cloning. We thank Michael Bonadonna, Beata Ramasz, Daniel Gimenes, and Diana Ordonez for helping with flow cytometry sorting. We thank Paola Grandi, James Reddington, Christine Rummel for critical feedback on establishing scifi-RNA-/scifi-ATAC-seq protocols. We thank the single-cell OpenLab of the Germany Cancer Research Center (DKFZ) and Jan-Phillip Malm for supporting with NGS runs. We thank Martin Garrido Rodriguez-Cordoba and Tim Pollex for critical feedback on this manuscript and Lars Steinmetz, Daniela Mauceri and the Noh laboratory for the helpful discussions.

## Author contributions

U.Y.: Conceptualization, Investigation, Methodology, Validation, Formal analysis, Writing - Original Draft + Review & Editing, Visualization. A.C.: Formal analysis, Resources, Data Curation, Writing - Review & Editing, Visualization. M.M.: Methodology, Formal analysis, Resources, Writing - Review & Editing, Visualization. A.C. and M.M. contributed equally. V.C.F.: Formal analysis, Resources, Data Curation. M.L.: Investigation, Formal analysis. M.Sar.: Formal analysis, Resources. M.Sav.: Formal analysis, Resources. D.B.: Investigation. M.W.D.: Supervision. J.B.Z.: Resources, Writing - Review & Editing, Funding acquisition, Supervision. K.M.N.: Conceptualization, Resources, Writing - Original Draft + Review & Editing, Funding acquisition, Supervision

## Data availability

Raw sequencing data will be uploaded on ENA and EGA. Accession numbers will be provided upon publication. All code used to generate intermediate files are available upon request.

## Ethics declaration

Human iPSCs used in this study were derived from peripheral blood mononuclear cells with institutional review board approval (Stanford University, reference numbers 29904, 30064) and approved by the EMBL Research Ethics Committee.

---

**Extended Data Fig. 1.**
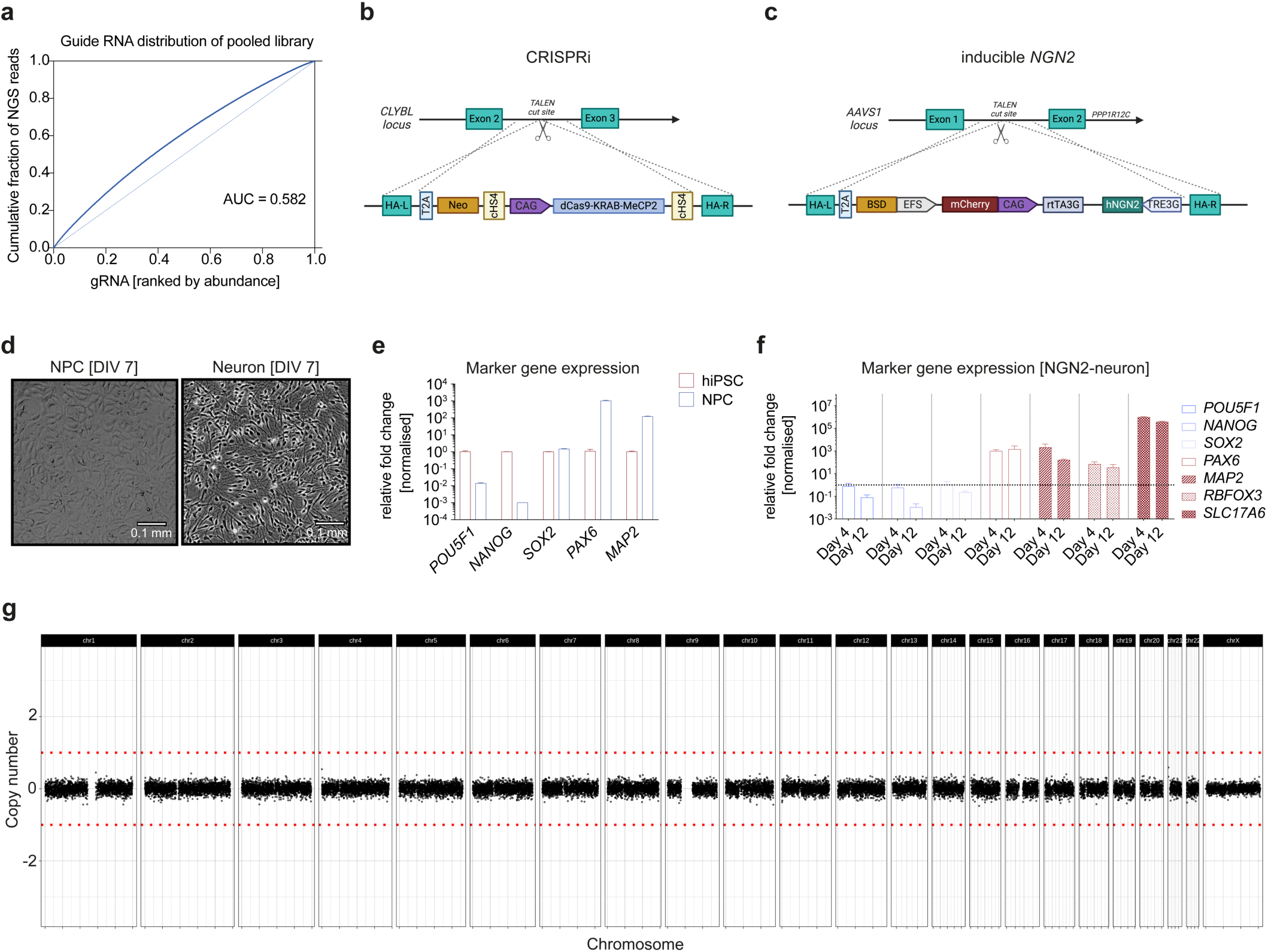
(**a**) Lorentz-plots showing the gRNA distribution after pooled cloning as a function of ranked gRNAs (x-axis) versus the cumulative fraction of assigned gRNA reads in relation to the library size (y-axis). The theoretic ideal distribution (AUC = 0.5) is indicated as a dotted line (light blue). *AUC: area under the curve*. (**b**) Strategy of introducing the CRISPRi machinery into the *CLYBL* safe- harbor locus via TALENs. *T2A: thosea asigna virus self-cleaving peptide motif; Neo: Neomycin resistance cassette; cHS4: chicken hypersensitive site 4 insulator; CAG: synthetic promoter, fusion of cytomegalovirus (CMV) enhancer to the chicken beta-actin promoter*. (c) Site-specific insertion of a doxycycline-inducible *NGN2* expression cassette into the *AAVS1* safe-harbor locus via TALENs. *BSD: blasticidin S deaminase resistance gene; EFS: EF1-*a *core promoter; TRE3G: tetracycline-response element-coding transgene; rtTA3G: tetracycline-inducible promoter (Tet-on).* **d**) Representative bright-field images of differentiating NPCs and neurons at day 7 post differentiation induction (DIV: days *in vitro*). (**e**, **f**) Expression of stem cell (*POU5F1*, *NANOG*, *SOX2*), progenitor (*SOX2*, *PAX6*) and neuronal (*MAP2*, *RBFOX3*, *SLC17A6*) markers in hiPSCs and derived NPCs (**e**) or in glutamatergic excitatory neurons following *NGN2* induction as measured via qPCR (**f**, day 4 and day 12 of differentiation exemplarily shown; ΔΔCt method; normalized to *GAPDH*/*ACTB* and experimental control (NT-ctrl). Error bars show mean ± the standard error of the mean (s.e.m; *n* = 4 independent replicates in (**e**) or *n* = 2 in (**f**)). (**g**) Whole-genome sequencing of engineered CRISPR-NGN2 hiPSC lines confirmed chromosomal integrity. The copy number of the whole genome of engineered clones is normalized to the parental cell line (NGN2-hiPSCs). Each dot represents a 100 kb window. Borders of allelic deletion or insertion are indicated with dashed lines (red). The plot shown here is a representative result for all established clonal lines.

**Extended Data Fig. 2.**
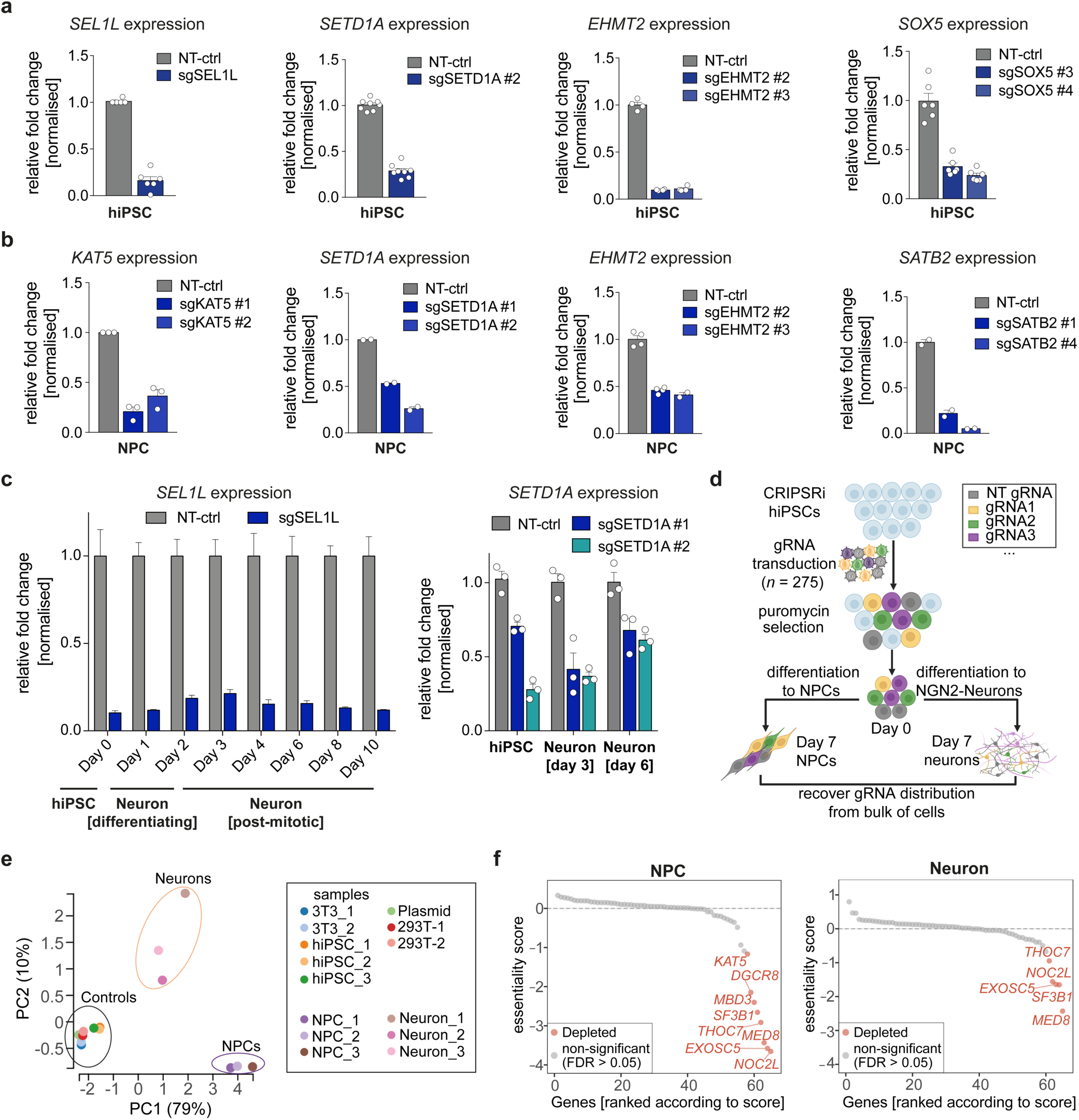
(**a-c**) Knockdown efficiencies of target genes in hiPSCs (**a**), NPCs (**b**), and throughout *NGN2*-driven differentiation (**c**) via CRISPRi (1 – 3 gRNAs for each target). Mean expression values were determined via qPCR (*n* = 1 – 8; indicated as individual data points when ≥ 2; ΔΔCt method; normalized to *GAPDH* and *ACTB* and experimental control (NT-ctrl)). Error bars show the mean ± s.e.m.. Target genes are indicated on top of the bar plots. (**d**) Schematic depicting the strategy of the bulk proliferation CRISPRi screen targeting schizophrenia risk genes. Pooled gRNAs (*n* = 275) are delivered to hiPSC constitutively expressing the CRISPRi machinery (*n* = 3 biological replicates) via lentiviruses at a low MOI (≤ 0.2). Cells are selected with puromycin treatment and allowed to recover. Four days post-transduction, cells are either harvested (to determine the baseline gRNA distribution; Day 0), maintained as hiPSCs, or differentiated to NCPs and induced neurons. Eleven days after pooled gRNA transduction (day 7 of differentiation) all cells are collected and gRNA distributions recovered via deep sequencing. (**e**) Principal component (PC) plot showing the first two principal components (PC1, x-axis; PC2, y-axis) calculated on gRNA count distributions. Percentage in parenthesis indicates the variance explained by each principal component. Samples of each group cluster together (groups are labelled on the plot). (**f**) Gene-level analysis of the screen in NPCs (left) and induced neurons (right). Essentiality scores (log2FoldChanges of normalized gRNA counts; y-axes) were calculated using MaGeCK^102^, normalizing gRNA counts of functional gRNAs to the NT-ctrl and to the gRNA distributions from day 0 hiPSCs. Genes are ranked and significantly depleted hits (FDR ≤ 0.05) highlighted in red.

**Extended Data Fig. 3.**
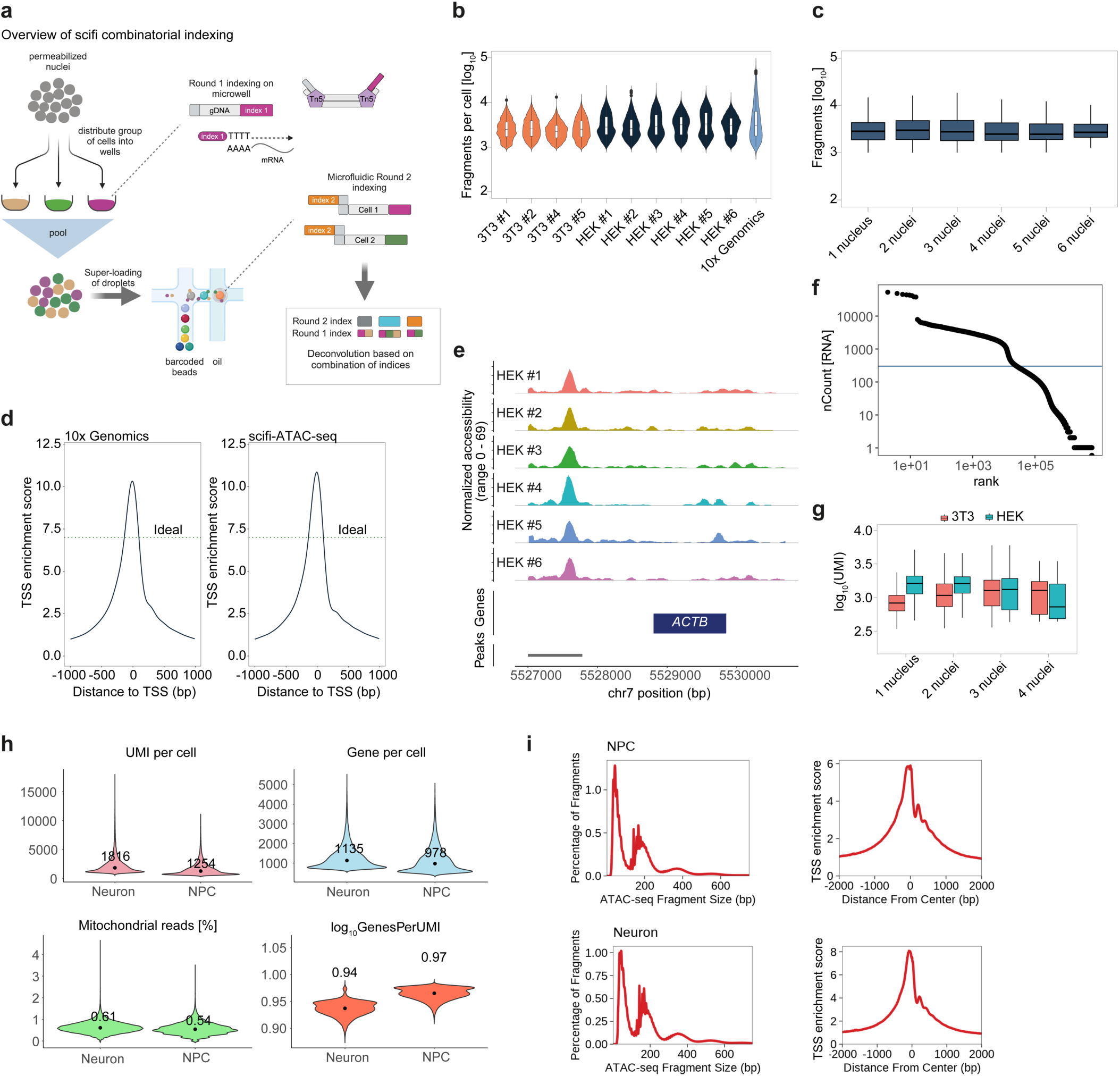
(**a**) Schematic of the combinatorial microfluidic barcoding for high-throughput scATAC- and scRNA-seq (scifi- ATAC-/scifi-RNA-seq). (**b-d**) Key QC metrics of the scifi-ATAC-seq assay: (**b**) fragments per cell (log10, y-axis) in each sub-library from the scifi-ATAC benchmarking (human samples in dark blue, mouse samples in orange). Data from a previously performed scATAC-seq run following the 10x Genomics Single-Cell ATAC kit was included for comparison (light blue). The median values are marked, and boxes indicate the 1.5x interquartile range. (**c**) Number of fragments (log10, y-axis) per nuclei encapsulated into singlets or multiplets (x-axis). (d) Transcription start site (TSS) enrichment comparison of the standard 10x Genomics scATAC- and the scifi-ATAC-seq readouts. The dashed line indicates an ideal sample according to ENCODE ATAC-seq standards (https://www.encodeproject.org/atac-seq/). (**e**) Aggregated chromatin accessibility peaks from HEK293T scifi-ATAC-seq readouts at the *ACTB* locus. (**f, g**) Key QC metrics of the scifi- RNA-seq assay: (**f**) Number of UMIs (y-axis) per cell barcode (x-axis). The line at y = 400 corresponds to the threshold value to separate true cells from ambient material and cells with compromised complexity. (**g)** Boxplots showing the UMI counts (y-axis, log10 transformed) for HEK293T (green) and NIH-3T3 (orange) across droplets that contained one or multiple nuclei (x-axis). (**h**) Quality control of scifi-RNA- seq in NPCs and induced neurons. Violin plots show the number of UMIs per cell (top-left), genes detected per cell (top-right), percentage of mitochondrial reads (bottom-left) and the estimated library complexity (log10(UMI per gene), bottom-right) in both cell types. The median is indicated on the violin plots. (**i**) Quality control of scifi-ATAC-seq in NPCs (top) and induced neurons (bottom). Left panels show the fragment size distributions of the sequencing libraries and the right panels the TSS enrichment scores (± 2 kilobases).

**Extended Data Fig. 4.**
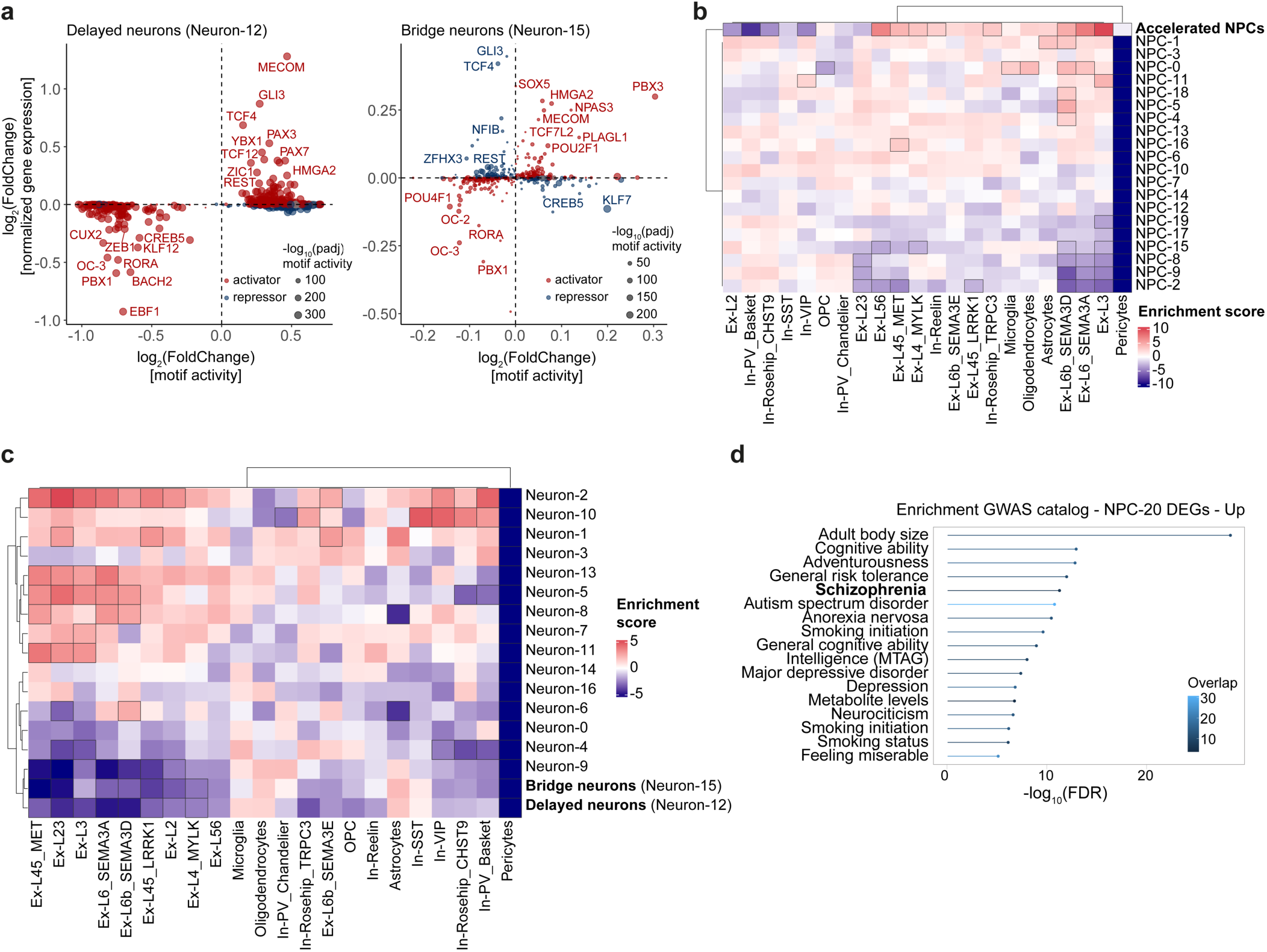
(**a**) Expression (y-axes) of TFs with differential activity (x-axes) in delayed (left panel) and bridge neurons (right panel). Differential activity and expression scores were computed contrasting against all other neuron clusters. TFs are labelled according to their inferred function as activators or repressors. Circle size indicates significance derived from motif enrichment analysis. Top hits are labelled. (**b, c**) Enrichment analysis using scDRS^105^ with DEGs in schizophrenia with an extended set of brain cell types in NPC (**b**) and Neuron clusters (**c**). Heatmaps show the scaled average enrichment for cell type specific schizophrenia DEGs. Brackets indicate significance (FDR ≤ 0.05). *Ex-L: excitatory layer; In: inhibitory; OPC: oligodendrocyte progenitor cell; PV: parvalbumin-positive; SST: somatostatin-positive; VIP: vasoactive intestinal polypeptide-expressing.* (**d**) Enrichment analysis of upregulated genes in accelerated NPCs (log2FoldChange ≥ 0.2; *n* = 268) with traits from the GWAS catalog.

**Extended Data Fig. 5.**
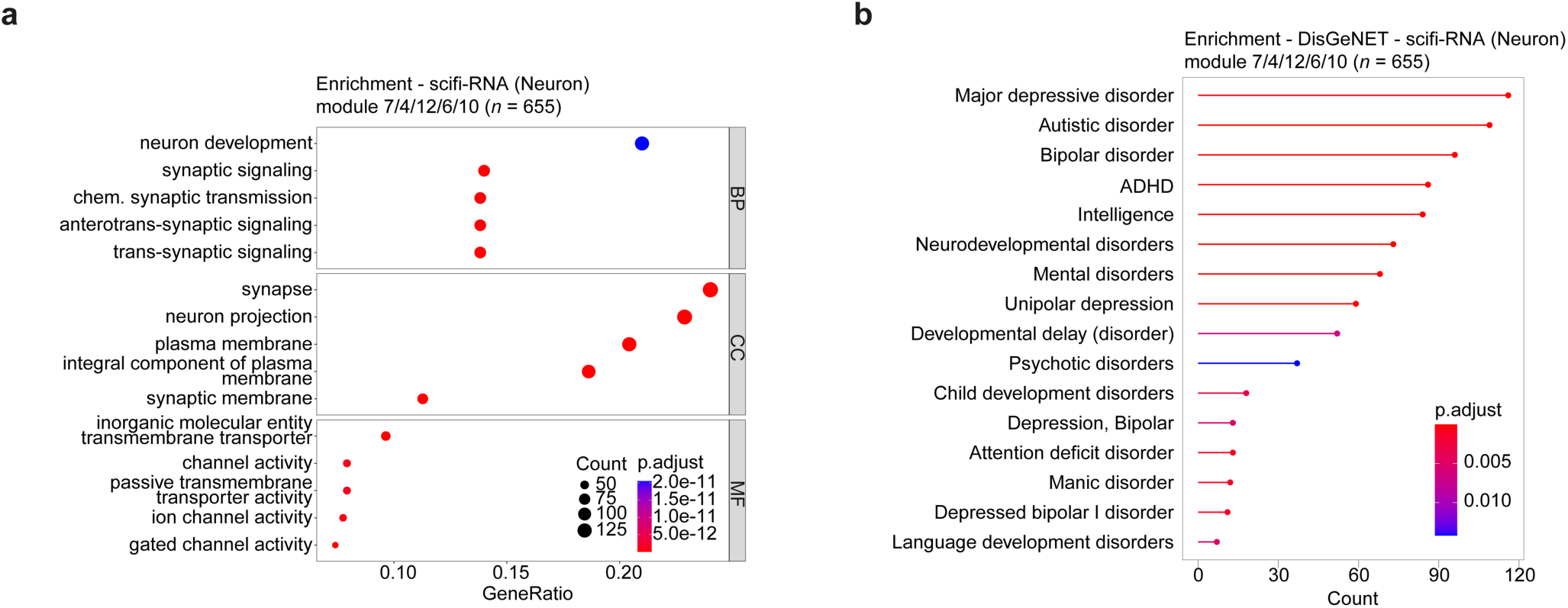
(**a, b**) Enrichment analysis of genes in modules 4, 6, 7, 10, and 12 (*n* = 655 genes) using Gene Ontology (GO) terms (**a**) or using disease terms from DisGeNET (**b**). Significant terms related to neurodevelopmental processes are shown. As background for the enrichment analysis, all variable genes were used that were employed for the construction of gene set modules (*n* = 3,216 genes).

**Extended Data Fig. 6.**
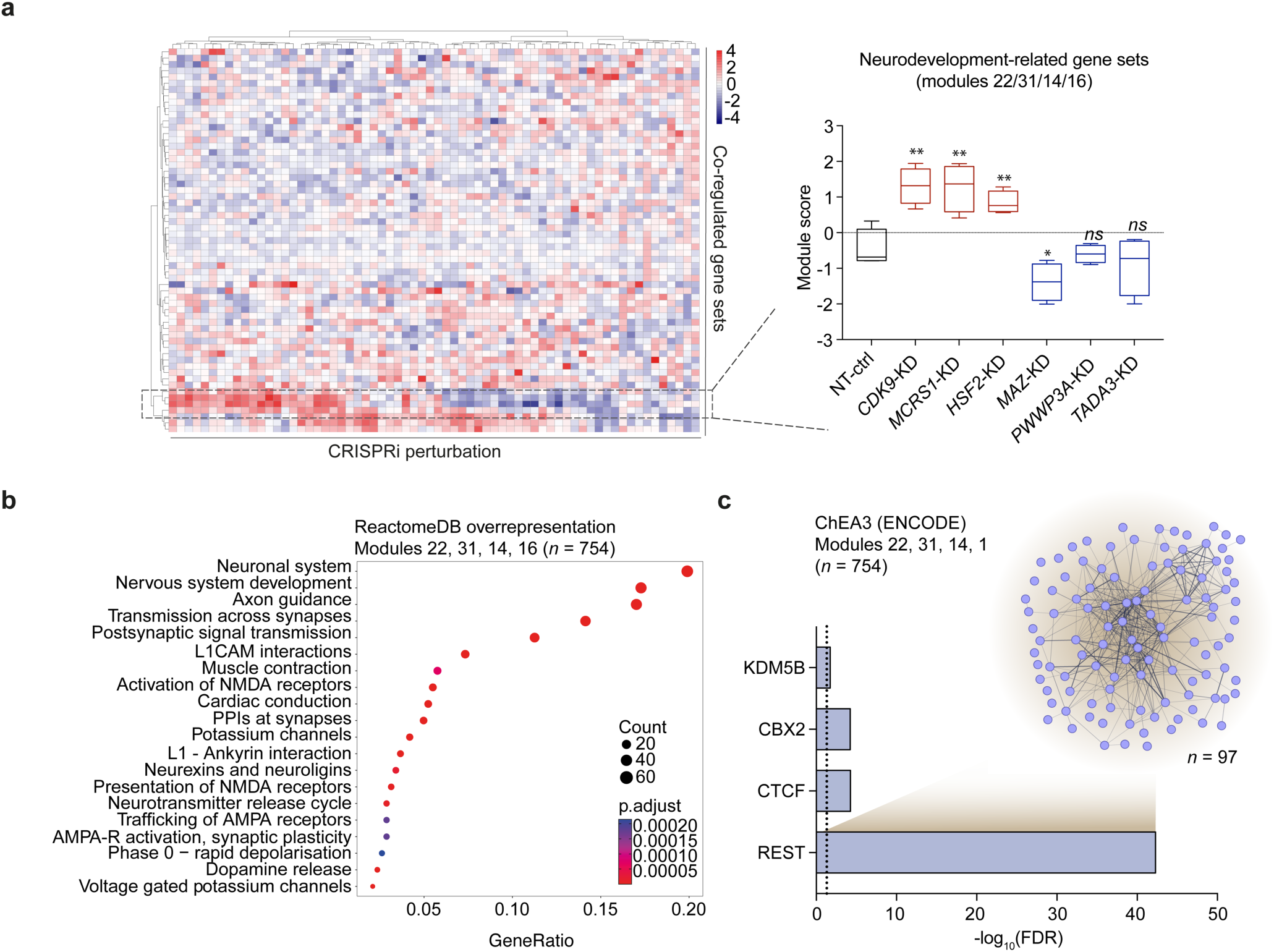
(**a**) Heatmap showing the expression of 61 gene set modules (indicated on the y-axis, generated using Monocle 3^58^) following CRISPRi (x-axis) in NPCs. Modules associated with neurodevelopmental processes (modules 14, 16, 22, 31; *n* = 754 genes) are highlighted with a dotted box. The average module score (modules 14, 16, 22, 31) contrasting the NT-ctrl group with CRISPR perturbation groups that showed an increased expression (*CDK9*-, *MCRS1-*, *HSF2-*KD) or reduced expression (*MAZ-*, *PWWP3A-*, *TADA3*-KD) shown as boxplots (center line represents the median; the lower and upper hinges represent the 25th and 75th quartiles, respectively; whiskers denote 1.5x of the interquartile range). Unpaired two-tailed t-tests were performed (* *P ≤* 0.05, ** *P* ≤ 0.01, *ns:* non- significant). (**b**, **c**) Enrichment analysis of neurodevelopmental gene sets with terms from the REACTOME database (**b**) or with TFs from ChEA3^103^. Dashed line in (**c**) indicates the significance threshold (FDR ≤ 0.05). Putative REST targets formed a STRING protein-protein- interaction network (confidence: 0.4, *n* = 97; disconnected nodes are hidden). All variable genes used for the gene module analysis (*n* = 8,664) was used as a background gene set for the enrichment analyses, while ChEA3 was run with standard parameters.

**Extended Data Fig. 7.**
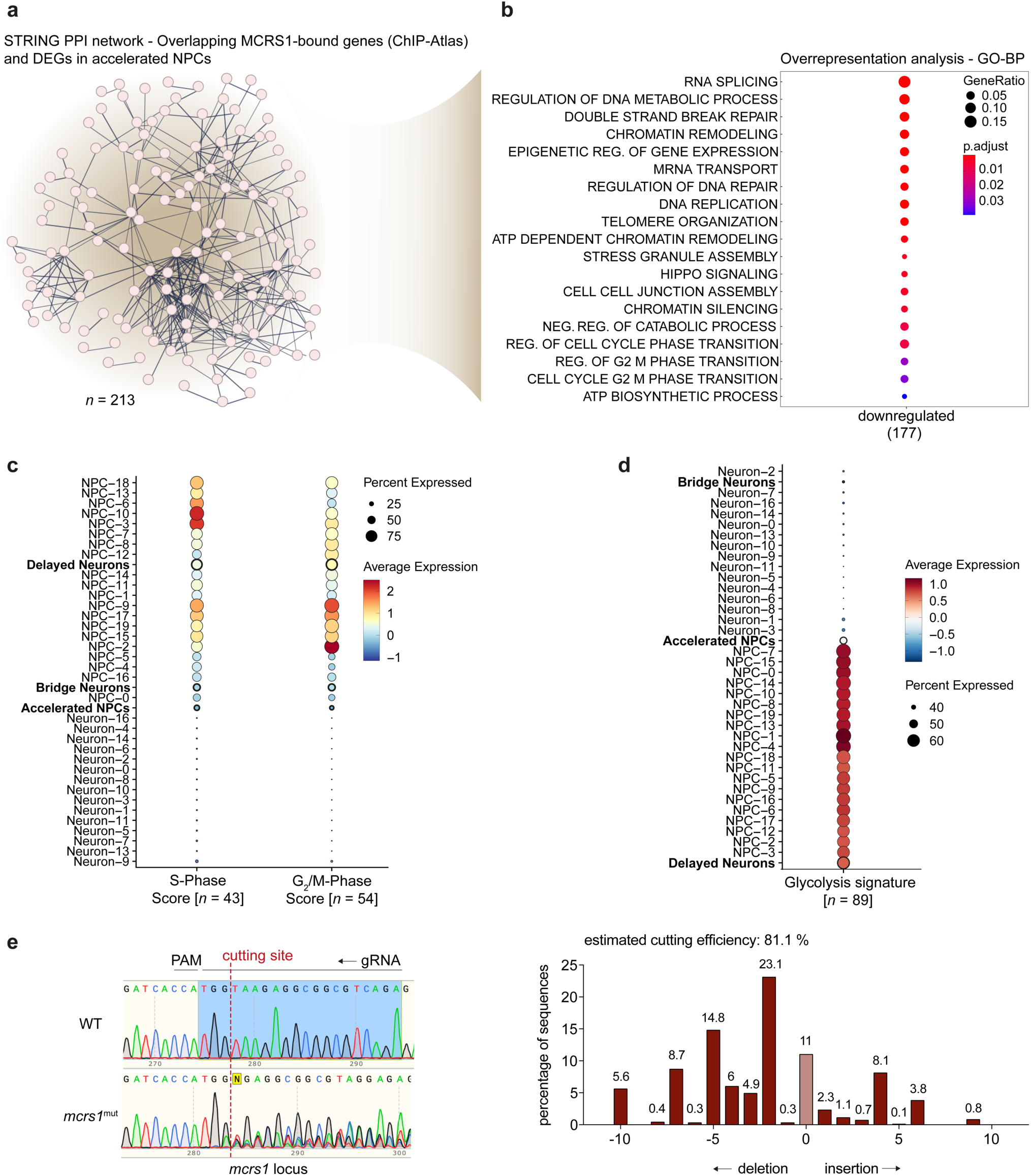
(**a**) Protein-protein-interaction network (STRINGdb, significance threshold: 0.7; *n* = 213; disconnected nodes hidden) of overlapping genes bound by MCRS1 (derived from ChIP-Atlas) and DEGs of accelerated NPCs (NPC-20 cluster). (**b**) Enrichment analysis of overlapping genes bound by MCRS1 and downregulated in accelerated NPCs (NPC-20 cluster) with GO-BP terms. Most significant terms for downregulated genes are shown (FDR ≤ 0.5). (**c**) Dot plot showing the mean expression of cell cycle marker (left S-phase, right G2/M; derived from^70^) in NPC and neuron clusters. Circle size indicates the percentage of cells expressing genes in the marker gene sets. Clusters of accelerated NPCs and delayed neurons (NPC-20, Neuron-12, and Neuron-15) and their scores are highlighted. (**d**) Dot plot showing the mean expression of glycolysis-related genes (retrieved from GO:0006096) in NPC and neuron clusters. The size of the circle indicates the percentage of cells expressing genes in the signature gene set. Clusters of accelerated NPCs and delayed neurons and their scores are highlighted. (**e**) Representative sanger traces of the *mcrs1* cut site in WT (top) and mutant (bottom) zebrafish (left panel) with one of the gRNAs. Cutting efficiency in mutant zebrafish was estimated using TIDE (right panel).

**Extended Data Fig. 8.**
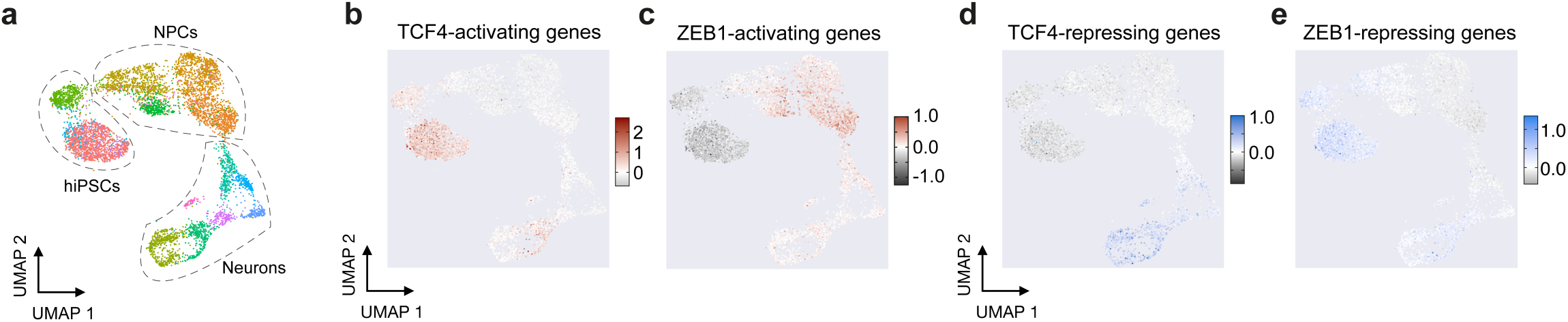
(**a**) UMAP showing the timecourse expression data of hiPSCs, NPCs and induced neurons. (**b-e**) UMAPs showing the results of the scDoRI analysis with the activation (**b**, **c**) or repression (**d**, **e**) of genes regulated by TCF4 (left) or ZEB1 (right).

**Extended Data Fig. 9.**
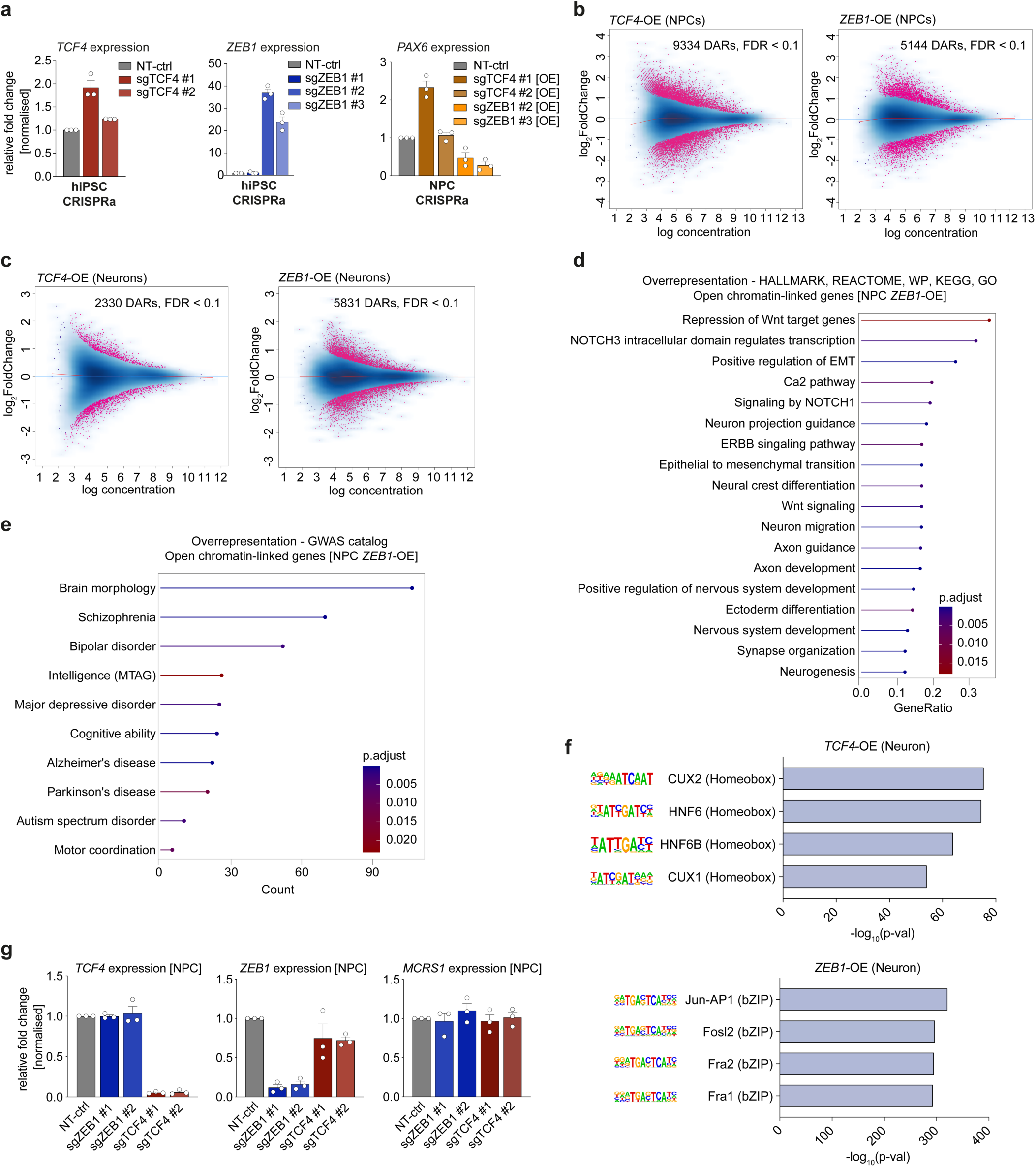
(**a**) relative expression of *TCF4* (left) or *ZEB1* (center), and *PAX6* (right) in control cells (NT-ctrl) and following CRISPRa of *TCF4* or *ZEB1* in hiPSCs (left and center panels) and NPCs (right). (**b, c**) Log ratio vs. average (MA) plots showing the significantly differentially accessible chromatin regions (DARs) following *TCF4-*/*ZEB1*-OE vs. controls in NPCs (**b**) and induced neurons (**c**). (**d**, **e**) Overrepresentation analysis of genes linked to differentially more accessible chromatin regions following CRISPRa of *ZEB1* in NPCs with ontology terms derived from MSigDB (Hallmark category), the KEGG, Reactome, Wikipathways (WP) and Gene Ontology (GO) databases (**d**) or with traits from the GWAS catalog (**e**). Significant terms are shown (FDR ≤ 0.05). (**f**) HOMER motif enrichment results in induced neurons following CRISPRa of *TCF4* (top) or *ZEB1* (bottom). Top four enriched TF motifs are shown with their significance (bar plot; -log10(p-val)). (**g**) Relative expression of *TCF4*, *ZEB1*, and *MCRS1* in control cells (NT-ctrl) and following CRISPRi of *TCF4* or *ZEB1* in NPCs. In (**a**) and (**g**), relative expression levels were measured via qPCR following the ΔΔCt method, normalized to *GAPDH* and *ACTB* and experimental control (NT-ctrl). Bar plots show mean ± s.e.m. (*n* = 3 independent replicates).

